# Ultra-Rapid Somatic Variant Detection via Real-Time Threshold Sequencing

**DOI:** 10.1101/2021.05.14.444172

**Authors:** Jack Wadden, Brandon Newell, Joshua Bugbee, Robert P. Dickson, Carl Koschmann, David Blaauw, Satish Narayanasamy, Reetuparna Das

## Abstract

Molecular markers are becoming increasingly important for cancer diagnosis, proper clinical trial enrollment, and even surgical decision making, motivating ultra-rapid, intraoperative variant detection. Sequencing-based detection is considered the gold standard approach, but typically takes hours to perform. In this work, we present Threshold Sequencing, a methodology for designing protocols for targeted variant detection on real-time sequencers with a minimal time to result. Threshold Sequencing analytically identifies a time-optimal threshold to stop target amplification and begin sequencing. To further reduce diagnostic time, we explore targeted Loop-mediated Isothermal Amplification (LAMP) and design a LAMP-specific bioinformatics tool—LAMPrey—to process sequenced LAMP product. LAMPrey’s concatemer aware alignment algorithm is designed to maximize recovery of diagnostically relevant information leading to a more rapid detection versus standard read alignment approaches. Coupled with time-optimized DNA extraction and library preparation, we demonstrate confirmation of a hot-spot mutation (250x support) from tumor tissue in less than 30 minutes.

## Introduction

Cancers are increasingly being diagnosed and treated based on underlying genetic driver mutations^1^ . Augmenting traditional histopathology with molecular diagnostics can not only improve diagnostic accuracy^2^, but also add prognostic value^3,4^ and inform disease management^5–7^ . If diagnosed intraoperatively, during biopsy or resection, certain molecular markers can lead to a change in surgical management^6,8,9^, combination of surgical procedures (e.g. biopsy and resection), and even targeted intraoperative treatment^10^ . This might even include intra-operative enrollment or exclusion from clinical trials that target certain mutation-specific pathways^5^ . As more actionable targets and targeted therapies are discovered, intraoperative molecular diagnostics will only increase in importance and utility, motivating the development of rapid, and easy to perform molecular assays.

In spite of their potential benefit, standard molecular diagnostics generally have long turn-around times (typically days to weeks) and cannot be practically performed within the intraoperative timeframe (which we define here to be <1hr based on several interviews with neurosurgeons who regularly take advantage of intraoperative diagnostics). Many traditional molecular diagnostic tests have been optimized to work intra-operatively with varying trade-offs (summarized in Supplementary Materials, Table S1). Immunohistochemistry (IHC) can detect certain biomarkers that correlate to actionable mutations^11,12^, but is difficult to extend and apply to other mutation hot spot loci or oncogenes. Similarly, Raman spectroscopy has been shown to enable rapid diagnosis of specific IDH mutations and 1p19q co-deletion^13^, but is currently limited in scope from diagnosing other clinically relevant targets. Intraoperative fluorescent in-situ hybridization (FISH)^14,15^ and targeted quantitative polymerase chain reaction (qPCR) assays^8,16,17^ have been shown to provide rapid, and specific detection of targeted mutations or other cancer biomarkers. However, both FISH and qPCR diagnostics also operate using a limited number (2-7) of allele specific, targeted primers and/or probes or other biomarkers and are not easily extended to commonly mutated oncogenes where many possible mutations (e.g. *KRAS p*.*G13, EGFR*) and loci^18^ (e.g. *TP53)* would be clinically relevant.

In contrast to single, allele-specific detection, genotyping via targeted amplicon sequencing can identify any potential point mutation and even small structural variants and copy number variations within a single amplicon. Sequenced reads also have the added benefit that they can be informatically inspected to identify spurious or off-target amplification or other potential issues, increasing confidence in positive, negative, and indeterminant results. Targeted amplicon sequencing can also identify other clinically relevant biomarkers such as copy number variations^19,20^, and Variant/Mutant Allele Fraction (VAF), which has increasing clinical relevance for prognostics, disease tracking, and management^21,22^ . Yet even the fastest next generation sequencing-based (NGS) tissue-to-diagnostic protocols take well over an hour to perform (summarized in Table S1) and are currently much too slow to be used during surgery. Furthermore, NGS sequencing is cost-optimized for sequencing of batched patient samples, and generally less accessible than other assay types, limiting the practical nature of intra-operative sequencing.

Oxford Nanopore Technologies (ONT) has developed a portable, low-cost sequencing device that provides a rapid library preparation protocol, and streaming, real-time access to sequenced DNA for immediate analysis. ONT sequencers capture and feed DNA strands through nanopores embedded in a membrane and measure ionic current disturbances across the pore. These current measurements correspond to base-pairs and can be fed to a basecaller algorithm to recover the specific DNA sequence (Figure 1a). ONT’s technology is capable of sequencing long DNA strands (>1Mbp) and is primarily marketed at use-cases where long reads are beneficial (e.g., de novo genome assembly and structural variant detection) but the technology’s rapid time-to-result and streaming output make it a promising candidate platform to explore the feasibility of intraoperative sequencing. Prior work using ONT sequencers performed targeted amplicon sequencing from pre-extracted DNA within ∼2.5 hours^23^, but an end-to-end sequencing-based diagnostic within the intraoperative time-frame has yet to be demonstrated.

**Figure 1.**
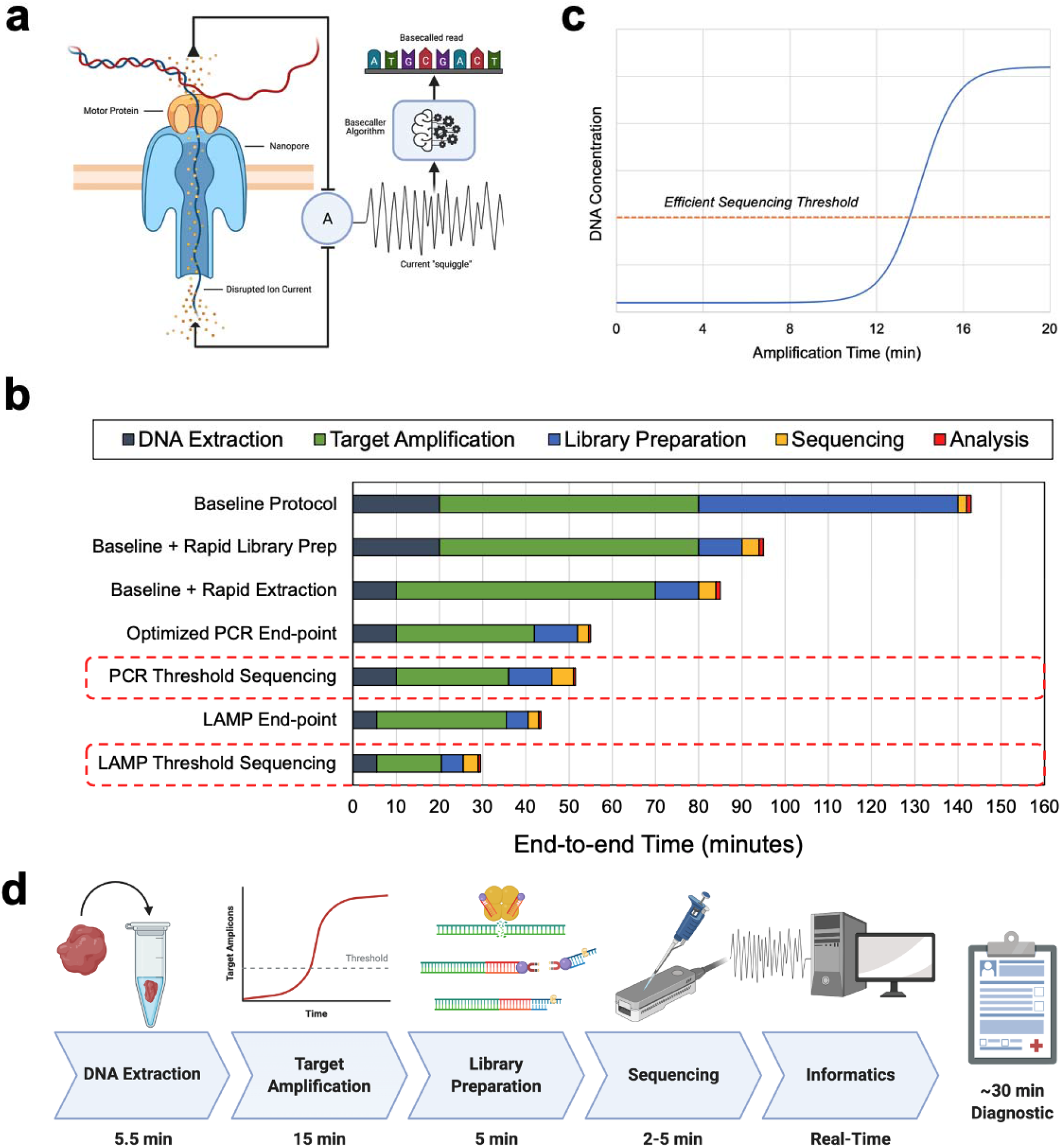
Overview of real-time Oxford Nanopore Sequencing, a time characterization of standard protocol and our optimization path, the intuition behind Threshold Sequencing, and our final end-to-end protocol. **(a)** Oxford Nanopore devices sequence DNA molecules by feeding a single strand through a pore embedded in membrane. Disturbances in electrical current across the membrane correspond to individual base pairs and can b reconstructed (basecalled) to form the DNA sequence. ONT sequences provide streaming sequencing output in real-time enabling immediate informatics and diagnostics (created with BioRender.com). **(b)** Characterization of time spent in various stages of ONT targeted sequencing pipelines. The baseline protocol is based off of standar suggested protocols for amplicon sequencing, and PCR cycling parameters used in prior work^23^ . Coupled with easy-to-perform rapid extraction and library preparation protocols, we demonstrate that sequencing-based diagnostics can be performed within the intraoperative timeframe. **(c)** Shallow target amplification allows enough amplification to provide diagnostically sufficient target depth and variant call support within a few minutes of sequencing, without wasting time on enrichment steps or overamplification. **(d)** Threshold sequencing, coupled with LAMP amplification, optimized library preparation, and optimized DNA extraction protocols can provide intraoperative sequencing-based diagnostic within 30 minutes (created with BioRender.com).

This work develops and demonstrates techniques for ultra-rapid (<1hr) end-to-end sequencing-based molecular diagnostics. Time characterization of protocols from prior work and standard ONT sequencing pipelines identified two major bottlenecks to achieving a <1hr diagnostic: library preparation and target amplification (Figure 1b). To reduce library preparation time, we investigate the suitability of ONT’s rapid 10-minute, fragmentation-based library preparation when applied to short (<300bp) amplicons. Short amplicons are not typically sequenced using this methodology due to concerns about over fragmentation, result quality, and a general lack of time-pressure. However, short amplicons are desirable for ultra-rapid assays because they generally amplify more efficiently leading to shorter amplification times. For this demonstration, we design primer sets covering known histone mutations—*HIST1H3B K27M* and later *H3F3A K27M*—present in previously characterized pediatric brain tumor tissue available in a local biorepository. We first evaluate the feasibility of the rapid library kit when applied to varying lengths of PCR amplicons and identify several trade-offs that impact time-to-result. We show that while fragmentation and the rapid library preparation do introduce various inefficiencies, these can be mitigated with design considerations, and are outweighed by the time benefit of the rapid approach.

To reduce target amplification time, we develop an amplification protocol optimization technique we call Threshold Sequencing. Rather than relying on a fixed, end-point protocol, Threshold Sequencing analytically identifies the optimal balance of target amplification time and amplicon sequencing time that leads to the fastest end-to-end result. Threshold sequencing works by first building a performance model of an ONT sequencer to estimate the sequencing time required to reach a desired level of diagnostic variant call support. The model can then be augmented with experimentally derived assay performance over time to predict the amount of target amplification time that leads to a time-optimal end-to-end diagnostic: the efficient sequencing threshold (Figure 1c).

To further reduce target amplification time, we investigate Loop-Mediated Isothermal Amplification^24^ (LAMP) as an alternative to PCR. LAMP uses isothermal strand-displacing polymerases, and intentional self-hybridization to rapidly generate and extend concatemeric amplicons. LAMP offers much more rapid target amplification than traditional PCR (∼30 minutes versus ∼1hr), making it an ideal candidate for rapid molecular diagnostics^25–29^ . However, LAMP assays can generate false positive amplification due to spurious mispriming events and primer self amplification^26,27,30^ . This spurious amplification is difficult to diagnose and negate without sequencing due to the many complex products formed that mimic or accompany proper amplification. Thus, sequencing and analysis of LAMP product is recommended to inspect and identify proper amplification^26^ . Current bioinformatics tools are not designed to process and diagnose complex concatemers (spurious or not) leading to difficulty in debugging assay performance, missed diagnostic information, and a slower diagnostic time-to-result.

We address these issues by designing an open-source LAMP concatemer analysis tool *LAMPrey* (https://www.github.com/jackwadden/lamprey).LAMPrey diagnoses and classifies each input read according to primer sequences and expected primer order generated by a properly behaved assay. LAMPrey thus allows for easy diagnosis of irregularities and inefficiencies in LAMP assays, including identification of suspected spurious amplification. Most importantly, LAMPrey can better identify diagnostically relevant target information than a standard long-read informatics pipeline. LAMPrey can also recover diagnostically relevant information lost to error-prone basecalling by aligning and polishing multiple redundant concatemer sections.

By time-optimizing DNA extraction and rapid library preparation protocols and leveraging rapid Loop-Mediated Isothermal Amplification (LAMP) amplification and the LAMPrey tool, we show that Threshold Sequencing can achieve 250x target support (mutant + wildtype calls) and successful somatic variant call from patient tumor tissue in less than 30 minutes (Figure 1b, 1d). This timeframe is comparable or better than the fastest targeted molecular diagnostics, while also offering the benefits of a sequencing-based approach. These feasibility experiments demonstrate that sequencing-based molecular diagnostics can be performed well within the intraoperative timeframe and have potential clinical utility.

## Results

### Threshold Sequencing Definition

Target amplification is performed primarily to increase the target signal over background genomic noise and typically involves conservative “end-point” protocols that maximize amplification (e.g. 30+ PCR cycles^23^). Long amplification protocols are generally beneficial, but at a certain point, the marginal benefit of more amplification does not contribute to a faster time-to-result. For a time-optimal diagnostic, amplification should only be performed if the time-to-result benefit outweighs the time cost of further amplification. Threshold Sequencing identifies this “worthwhile” threshold for amplification time and provides protocols that optimize for time-to-result.

For this definition, we assume ONT sequencers sample DNA strands from an input library uniformly at random. Therefore, amplicons will be sequenced/sampled according to the proportion of amplicon product relative to starting genomic DNA used as input template. More amplification time will increase total double stranded DNA mass, as well as the proportion of target amplicons relative to background genomic noise. Importantly, once sequenced on an ONT device, a DNA read is immediately available for downstream analysis, enabling real-time, streaming diagnostics, rather than a batched approach typical of other NGS technologies^31^ .

As an illustrative example of threshold sequencing, consider perfectly efficient PCR-based amplification where the target amplicon mass doubles every cycle. Given any amount of input DNA from any number of PCR cycles, we can model total diagnostic time using the formulas shown in Figure 2a. As amplicon mass grows, the proportion of total mass that is background genomic material (gDNA) reduces (Figure 2b) and thus the sequencing rate of target amplicons (versus background genomic DNA) increases. Assuming a 250x target coverage requirement for diagnosis, at a certain point, it becomes a waste of time to continue amplification, and instead becomes worthwhile to begin sequencing. Figure 2c shows an illustration of this trade-off using optimistic sequencing and PCR cycling parameters and a more sophisticated model of sequencing time (Supplementary Material Table S4). Assuming perfect PCR efficiency, the time-optimal PCR cycle threshold is 16 cycles, before further amplification results in wasted diagnostic time. This represents a savings of ∼12-22 minutes (15-20 cycles) versus a typical 30-35 cycle end-point PCR-protocol.

**Figure 2.**
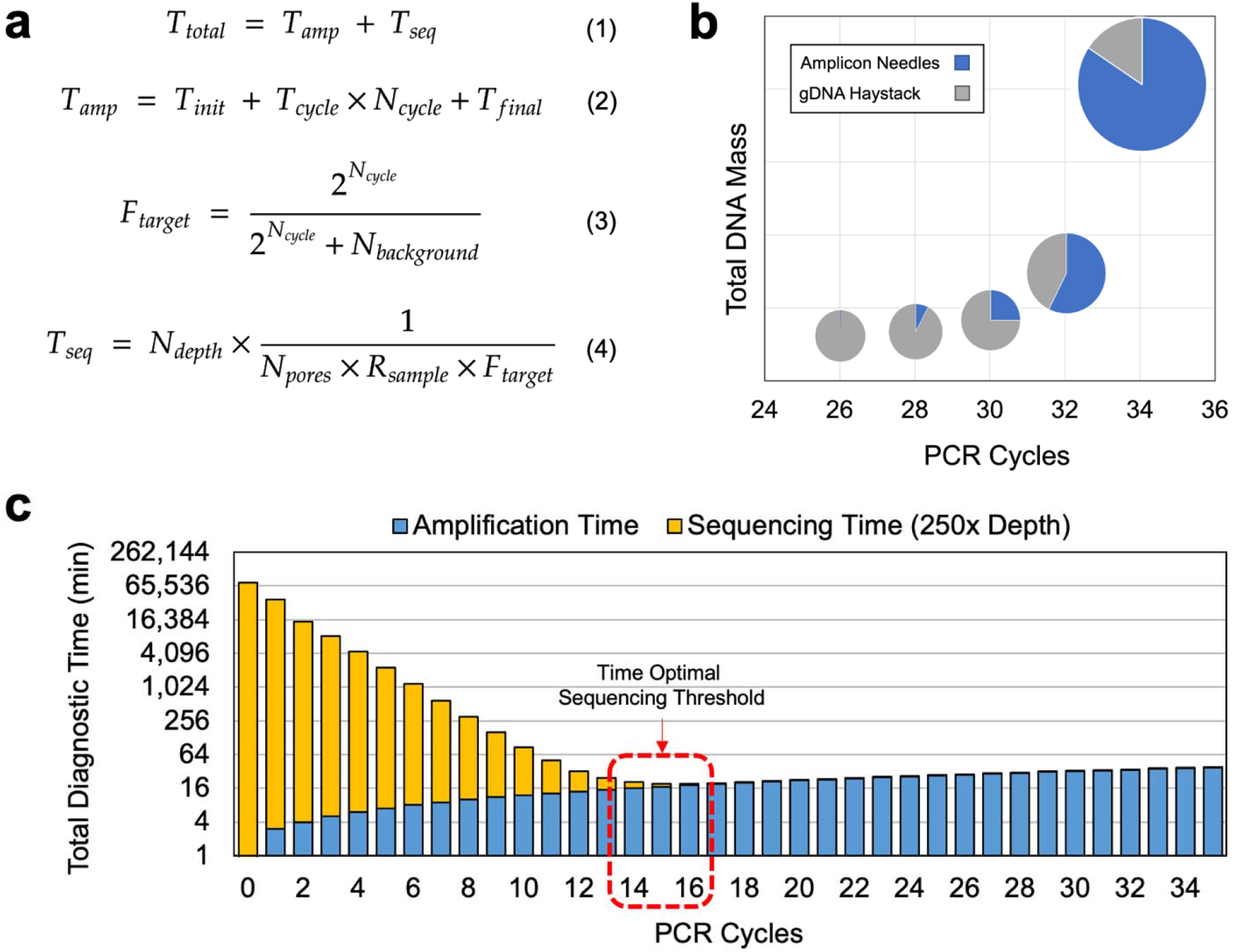
Threshold Sequencing identifies the amount of amplification time or cycles required to minimize total diagnostic time. **(a)** Diagnostic time can be modeled as the sum of amplification time (*T*_*amp*_) time required to sequence to a target diagnostic depth. Amplification time (*T*_*amp*_) is a function of cycling parameters in the case of PCR or incubation time in the case of LAMP. Sequencing time is a function of the target depth requirement (*N*_*depth*_), available parallel sequencing pores (*N*_*pores*_), the read sampling rate per pore (*R*_*sample*_), and the fraction of all sampled reads that are diagnostically useful (*F*_*target*_). **(b)** Over time, both total mass and proportion of amplicons relative t background genomic template (*F*_*target*_) increases. As amplicon “needles” increase, less sequencing time is required to search for them in the background genomic DNA (gDNA) “haystack.” **(c)** Using example PCR parameters, we can model total diagnostic time, and identify an amplification threshold that results in a minimal time-to-result.

In practice, it is difficult to predict amplification rate over time, especially considering LAMP amplification. Furthermore, other inefficiencies such as spurious LAMP amplification and amplicon fragmentation during library preparation will greatly impact the resulting useful target amplicon fraction (*F*_*target*_), sequencing rate, and time-to-result. To perform Threshold Sequencing in practice, we first identify useful target amplicon fraction and sequencing rate experimentally for a particular assay and tissue type, and then use the Threshold Sequencing model to identify a range of amplification times that provide required target mutation coverage for a particular use-case with an efficient use of time.

### Rapid Library Preparation and Feasibility and Optimization

Rapid library preparation time is key to enabling an intra-operative sequencing-based diagnostic. The recommended library preparation for amplicon-based sequencing on ONT platforms is ONT’s ligation sequencing kit (ONT #SQK-LSK109/110). However, this protocol takes between 40-60 minutes to perform and is assumed to be unsuitable for an intraoperative diagnostic. Oxford Nanopore also offers a rapid library preparation chemistry^32^ (ONT #SQK-RAD004) that is advertised as a 2-step, 10-minute protocol. In the first step, DNA is simultaneously fragmented and tagged with “click chemistry” (tagmentation) via a transposome complex (5 minutes). In the second step, tagmented, double-stranded DNA is mixed with click-chemistry prepared ONT sequencing adapters and allowed to incubate at room temperature for adapter attachment (5 minutes). This protocol is rapid and easy to perform but is not recommended for use with short amplicons. This is most likely due to concerns over transposome fragmentation behavior on short DNA fragments,^33,34^ which might be inefficient and generate extremely short, unmappable fragments. Regardless, short amplicons are desirable for targeted diagnostics because they generally allow for more efficient and rapid amplification.

To evaluate the feasibility of using the rapid library preparation kit on short amplicons we designed, amplified, and sequenced various lengths of tailed PCR amplicons (187bp, 260bp, 609bp, 910bp) using the rapid library kit. Primers were designed to cover the *HIST1H3B K27M* hotspot mutation present in pediatric brain tumor samples available to us via a biorepository (see Methods section). *HIST1H3B K27M* and *H3F3A K27M* are recurrent hotspot mutations in pediatric high-grade gliomas and predict especially poor prognosis.

We were successfully able to sequence and align all amplicons using standard bioinformatics pipelines (see Methods section), indicating that ONT’s rapid transposome-based library preparation is capable of tagmenting short amplicons down to 187bp in length. However, various inefficiencies were noted (Figure 3a). Some sequenced reads were not able to be successfully mapped to the human genome. This loss was between 8%-20% (Mapping Loss; Figure 3b) among all amplicon sizes and decreased as the amplicon length increased. This indicates that longer amplicons tend to fragment into longer reads which are generally easier to map.

**Figure 3:**
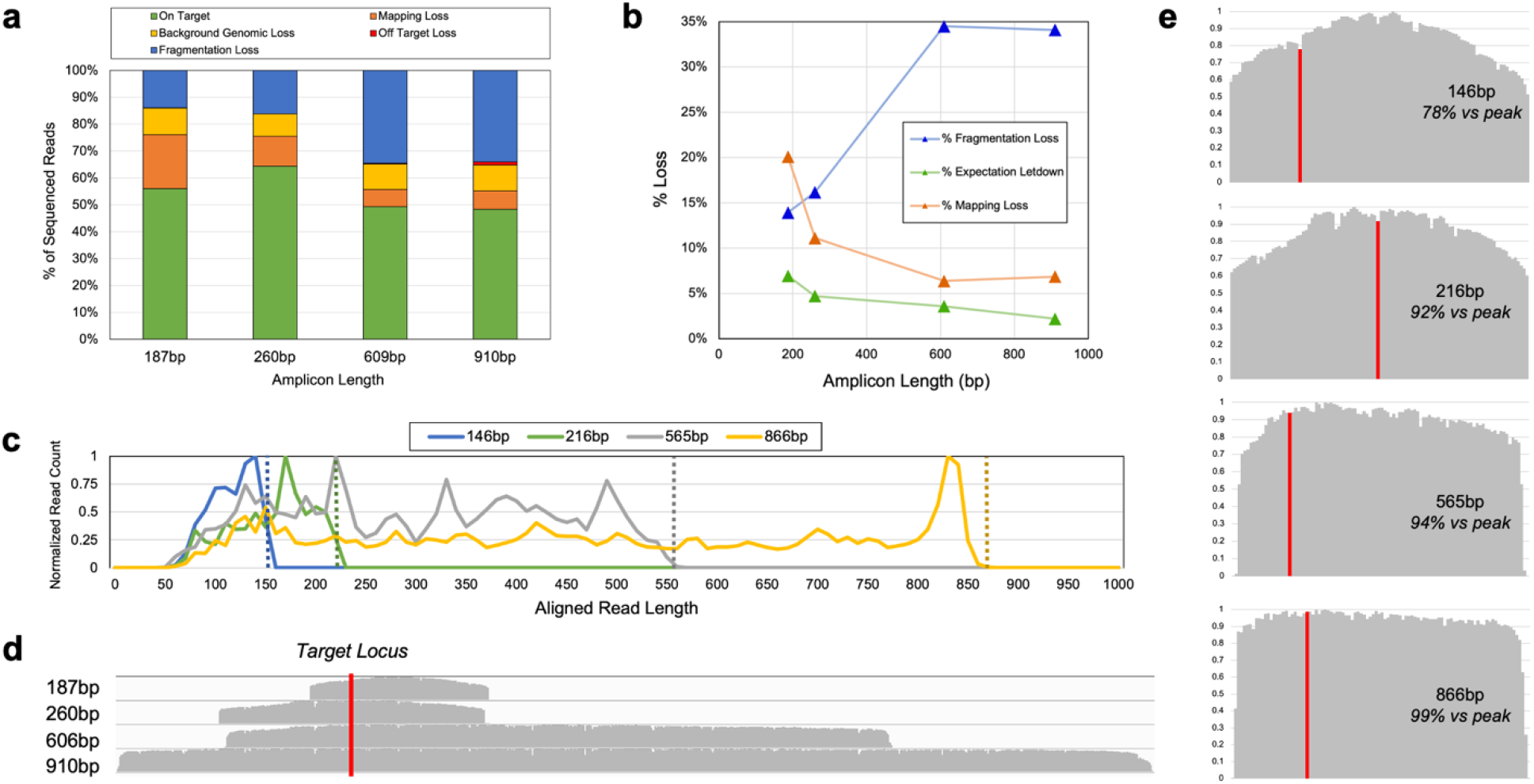
Fragmentation-based library preparation and its behavior when sequencing short amplicon products. **(a)** Due to the presence of background genomic DNA, fragmentation-based library preparation, and difficulty mapping short amplicon fragments, many sequenced reads are not diagnostically relevant, leading to additional sequencing time required to reach a desired target depth. **(b)** Longer amplicons tend to fragment int longer reads, which are easier to map (Mapping Loss). Longer amplicons and genomic DNA also tend to be favored by the transposome when compared to shorter amplicons (Expectation Loss). However, longer amplicons can fragment multiple times, producing a higher proportion of read fragments that do not contain a particular target, hot spot locus (Fragmentation Loss). **(c)** Aligned read lengths for all amplicons. Shorter amplicons lead to a larger proportion of reads that are too short to align properly. **(d)** Aligned read coverage histograms for all amplicons. **(e)** Width normalized coverage histograms for each amplicon showing edge penalties when the location of interest is near close to the end of amplicons. These results indicate it is beneficial to design primer sets that generate relatively short amplicons with any hot-spot target of interest near the center of the amplicon (to reduce fragmentation loss), but long enough to preserve the proportion of useful, mappable reads.

Based on the final amplified mass relative to the input template genomic DNA (measured via Qubit, Invitrogen dsDNA HS Assay #Q33230), we also would expect amplicons to be sequenced in roughly the same proportion. However, we observed a bias away from this expectation in favor of background genomic DNA (Expectation Letdown; Figure 3b). This indicates the transposome favors interacting with, and fragmenting longer genomic reads and longer amplicons versus shorter amplicons^34^ . While a trend favoring longer amplicons exists, even the shortest amplicon (187bp) incurred a loss of only 6.9% (versus 2.2% for the 910bp product) relative to amplified mass, an acceptable loss given the advantages of shorter amplicons and the rapid library preparation.

Due to the fragmentation involved in the rapid library preparation, a sequenced amplicon fragment might not contain a specific target locus of interest. Fragmentation loss (Fragmentation Loss; Figure 3b)—defined here as the reads that map to the target region but do not cover the target locus—is notably worse for longer amplicons and led to significant efficiency losses versus shorter products. This is most likely due to multiple transposomes fragmenting longer amplicons in multiple locations and producing a higher ratio of strands in the library that do not contain the target locus. The transposome will also fragment amplicons near their ends, producing both nearly full-length amplicons, and fragments that are too short to map properly map. This leads to an uneven distribution of aligned read lengths (Figure 3c)—especially for shorter amplicons—with sequencing and informatics favoring longer strands (Figure 3c), producing higher coverage near the middle of amplicons (Figure 3d/e).

It is therefore desirable to design primer sets such that hot-spot targets of interest lie as close to the center of the amplicon as possible to help reduce fragmentation loss. Even considering this loss, our results show that ONTs rapid, fragmentation-based library preparation kit is compatible with shorter amplicon lengths that allow for more rapid PCR cycling parameters. Given the relative performance of the 216bp (260bp with tails) primer set, we chose this pair for final cycle parameter optimization and end-to-end diagnostic evaluation. Final PCR cycle parameter optimization is described in Supplementary Materials.

To reduce total library preparation time, we explored reducing the suggested rapid adapter incubation time and its impacts on sequencing performance (Supplementary Material Figure S1). For four different time-points (2-5 minutes) we identified no clear trend in relative sequencing performance within the first 10 minutes of sequencing (Figure 3d). This indicates that a 2-minute incubation is at least sufficient to generate a sequencing library of acceptable quality for the ultra-rapid use-case, and further reduces the library preparation time by 3 minutes. This shortened protocol was leveraged in the final optimized intra-operative LAMP end-to-end run.

### Rapid DNA Extraction Evaluation

Rapid DNA extraction is also key to enabling a practical intra-operative diagnostic. Typical high-quality DNA extraction, isolation, and purification kits involve tissue digestion and multiple washing and elution steps with special binding columns^35^ . In practice, these centrifugation protocols tended to take longer than advertised (>20 minutes) due to multiple hands-on steps motivating the evaluation of more rapid approaches. We chose to evaluate a rapid, 8-minute extraction kit—Lucigen QuickExtract^36^ (LQE)—which employs a “one-pot” protocol, can use a raw piece of biopsied tissue as input, requires no centrifugation or isolation steps, and generates DNA immediately suitable for input to PCR amplification and LAMP. The LQE protocol involves a 6-minute extraction incubation and a follow-up 2-minute enzyme de-activation incubation. While rapid, the resulting DNA is relatively impure and requires dilution to achieve efficient amplification. To explore the feasibility of using LQE for our ultra-rapid exploration, we first followed the standard 8-minute protocol and confirmed that with physical overheads such as intermittent vortexing, pipetting, and handling, the protocol was easy to perform and took around 10 minutes in practice.

To measure the suitability of this DNA product for use in downstream PCR and sequencing, we also explored PCR efficiency of various dilutions of LQE product DNA in nuclease free water. We performed serial dilutions of the same LQE product, performed PCR, and confirmed specificity and efficiency via gel electrophoresis (Supplementary Material Figure S2). The most efficient PCR reactions were between 1:10 and 1:20 dilution ratios. We thus recommend a 1:10 dilution of LQE product as a baseline input to PCR to maximize product and total DNA as input to sequencing. We also verified that this PCR product could be sequenced using the ONT Rapid Library chemistry and MinION flow cell (Supplementary Material Figure S3) without unacceptable degradation of the flow cell membrane. Sequencing was able to be performed successfully without any modifications to the standard rapid library preparation and sequencing protocols.

Further time reduction was achieved by shortening the DNA extraction incubation time from 6 minutes to 1 minute and replacing the intermittent vortexing with a single initial 30s vortex in a smaller volume of LQE (100ul) to quickly disrupt tumor tissue. This technique led to final product with less total DNA, but with almost identical amplification rate to DNA extracted using the full protocol (Supplementary Material Figure S4). The shortened protocol was leveraged in the final threshold sequencing evaluation and optimized intra-operative LAMP protocol.

### PCR-based Threshold Sequencing

A final Threshold Sequencing amplification protocol can be designed by minimizing the diagnostic time Equation 1 and 4 from Figure 2a using experimentally derived values for expected amplification time (*T*_*amp*_), target fraction (*F*_*target*_), read sampling rate (*R*_*sample*_), and the number of pores available for sequencing on a MinION flow cell (*N*_*pores*_). Target fractions (*F*_*target*_) for various cycles of PCR were determined by performing our optimized PCR protocol for varying numbers of cycles. We then measured both the resulting PCR product concentration via Qubit fluorometer and sequenced each resulting product using a barcoded, multiplexed rapid library preparation methodology. Resulting reads were aligned to the human reference and classified according to alignment (see Methods section). Over time, we see both the total mass, as well as the proportion of PCR product relative to background genomic reads grow (Figure 4a).

**Figure 4.**
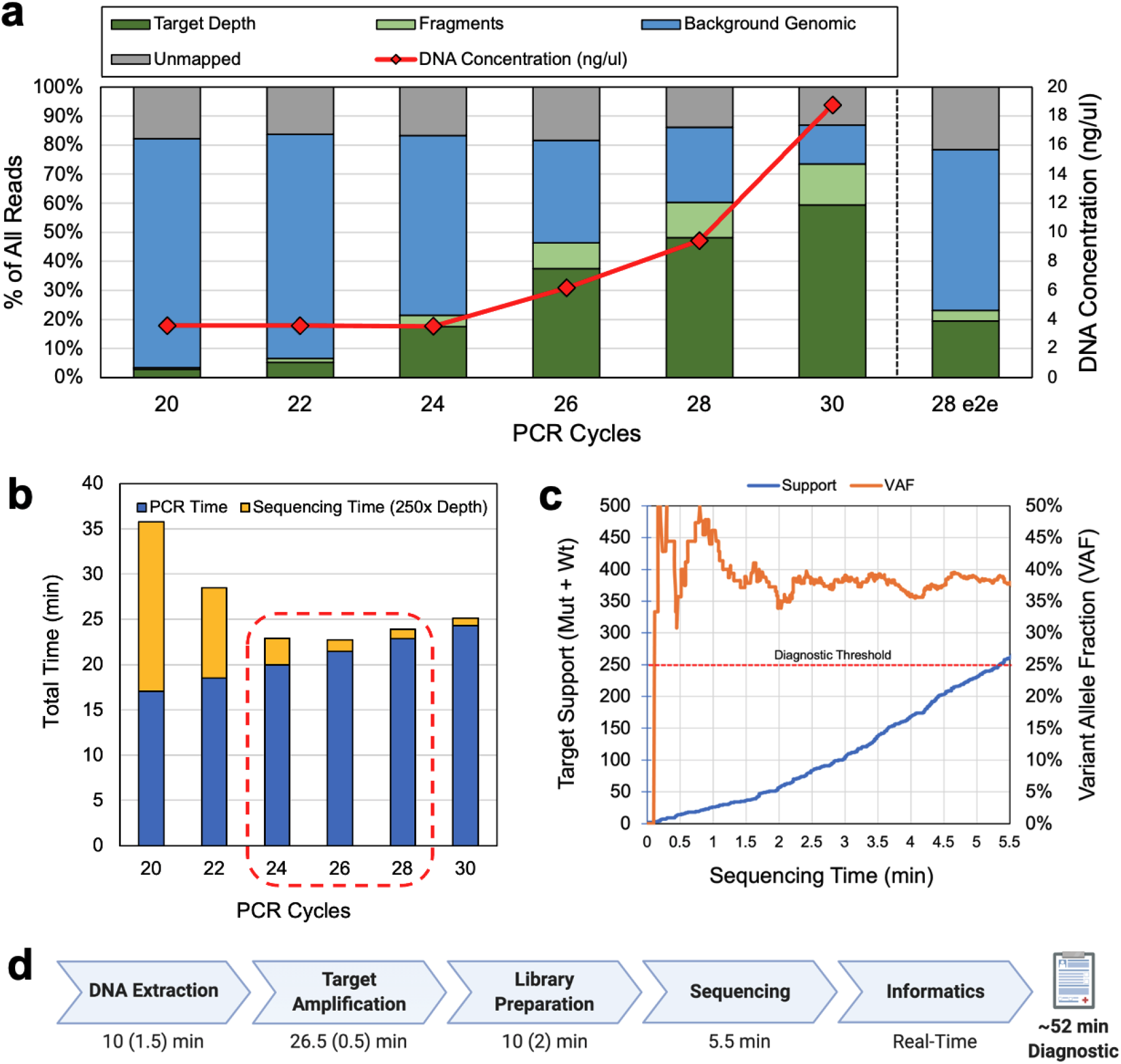
Threshold sequencing PCR cycle estimation and end-to-end run results. **(a)** Sequencing results and DNA concentration from various cycle counts of PCR product and the final end-to-end demonstration (28 e2e). As mass increases, corresponding target fraction (*F*_*target*_) increases. **(b)** Threshold sequencing model estimates corresponding to target fraction recovered from PCR cycle sweep identify optimal diagnostic time between 24 and 28 cycles. Overprovisioning PCR cycles offers less of a time-penalty than underprovisioning due to the exponential nature of PCR signal amplification and linear nature of ONT DNA sequencing. **(c)** In and end-to-end demonstration of the final protocol, using 28 cycles of PCR, we were able to reach the estimated diagnostic within 5.5 minutes of sequencing time. **(d)** Approximate times for each diagnostic step with approximate manual overhead in parenthesis (Created with BioRender.com). Informatics was performed after the fact and verified to be performable in real-time with negligible overheads resulting in a ∼52-minute diagnostic.

ONT read sampling rate (*R*_*sample*_) was estimated by measuring sample-rate from representative sequencing runs. Sequencing rate was measured to be 5.06 reads/s or 0.0158 reads/channel/s by extracting read channel and timestamp metadata (included in FASTQ output by the Guppy basecaller) using a custom script (see Code Availability). This measured rate was relatively low compared to prior results from standard libraries prepared from higher concentrations of high-molecular weight DNA, and most likely due to low input DNA concentration caused by extraction dilutions, leftover inhibitory reagent, and the omission of a PCR product cleanup step. Typical measured input DNA to library preparation was ∼20ng-60ng, much lower than that recommended by ONT (i.e. 400ng^32^ of high molecular weight DNA). However, we found DNA concentration difficult to increase without adding unacceptable amounts of time to the end-to-end diagnostic.

Assuming a target coverage requirement of 250x (*N*_*depth*_), 512 active channels available on a MinION flow cell (*N*_*pores*_), and our estimated *R*_*sample*_ and *F*_*target*_, total assay time is expected to be minimal between 24 and 28 cycles (Figure 4b). 250x coverage was chosen as a semi-arbitrary threshold to allow for highly confident calls from common heterozygous somatic mutant allele fractions found from patient tumor samples (>10%). In reality, depth requirement for statistical significance depends on the underlying sample type (e.g., core biopsy vs solid tumor), variant allele fraction, desired confidence level, and error rate expected from a particular basecaller algorithm. We discuss further motivation for picking 250x as a conservative diagnostic target and statistical methods for on-line variant calling in Supplementary Materials.

To demonstrate the time benefits of threshold sequencing, we performed a timed end-to-end run of the protocol starting with acquisition of ∼20mg of room temperature tumor tissue, performing rapid DNA extraction, 28 cycles of PCR, ONTs rapid library preparation, and sequencing (see Methods Section) and analysis to 250x target support (mutant + wildtype calls over the target). Because additional sequencing time penalty due to under-amplification is more severe than the time cost of extra PCR cycles (due to the exponential nature of PCR, and cycle-time optimization) we chose 28 cycles to account for unexpected inefficiencies and safeguard a result that fits within the intraoperative timeframe. Target support over time and sample variant allele fraction is shown in Figure 4c. Our protocol (see Methods section) was able to achieve 250x support and successfully call a known *HIST1H3B K27M* variant from a pediatric DIPG tumor sample in ∼52 minutes. Data analysis was performed after the fact but verified to be performable in real-time. The protocol is shown in Supplementary Material (Figure S7) and available online (see Methods Section). The approximate breakdown of time taken in each protocol step, including human overhead, is shown in Figure 4d.

Sequencing time overshot model estimates by ∼4 minutes. This was mostly due to a lower than expected *F*_*target*_ (19%) versus predicted (∼50%) increasing sequencing time by 3.6 minutes. This could be due to unexpected PCR inefficiency, or perhaps accidental over-fragmentation during the rapid library preparation leading to loss of small amplicon fragments and an increased proportion of genomic fragments. The ONT flow cell used for evaluation also was only able to activate 443 channels out of the 512 on the device, and only 409 channels/pores were able to participate in sequencing within the first five minutes (a 20% penalty versus model predictions). However, the read sampling rate per available channel was ∼1.8x higher than predicted somewhat compensating for other inefficiencies.

Further reductions to target amplification time were explored, including a 2-step PCR protocol. We also explored ONTs four-primer protocol that attaches click chemistry via a secondary PCR step with a second set of primers that include click chemistry (obviating fragmentation and eliminating the fragmentation penalty). Their evaluation is provided in Supplementary Material.

### LAMP Optimization and Efficiency Evaluation

While PCR-based threshold sequencing can provide a sequencing-based diagnostic within the intra-operative timeframe, PCR accounts for ∼50% of end-to-end assay time and is the largest performance bottleneck (Figure 1b). Loop-mediated Isothermal Amplification^24^ (LAMP) offers more rapid target amplification than PCR (generating micrograms of product within tens of minutes) and was chosen as a candidate to further reduce target amplification time.

LAMP operates by carefully choosing a template region and primer sequences such that product intentionally forms self-hybridizing loops. LAMP initiates with hybridization and extension of tailed “inner-primers”. This initial sequence is then displaced from the template via strand invasion from an “outer primer”. Inner primers are tailed with reverse complementary sequences designed to self-hybridize with the template, forming hairpin sequences, and eventually “dumbbell” seed structures. Once dumbbells are formed, extension can initiate from 3’ looped ends or inner-primer annealing sites on the loops themselves resulting in rapid concatemeric extension and copying of the target region (Figure 5a). LAMP amplification has been used in a variety of use-cases where rapid time-to-result is important including pathogen detection^26,30^ and targeted-allele genotyping^29,37^ . However, the complex machinery of LAMP can easily malfunction and lead to spurious amplification. By sequencing and properly analyzing LAMP product, we can diagnose issues with LAMP assays, and create both fast, and reliable assays.

**Figure 5.**
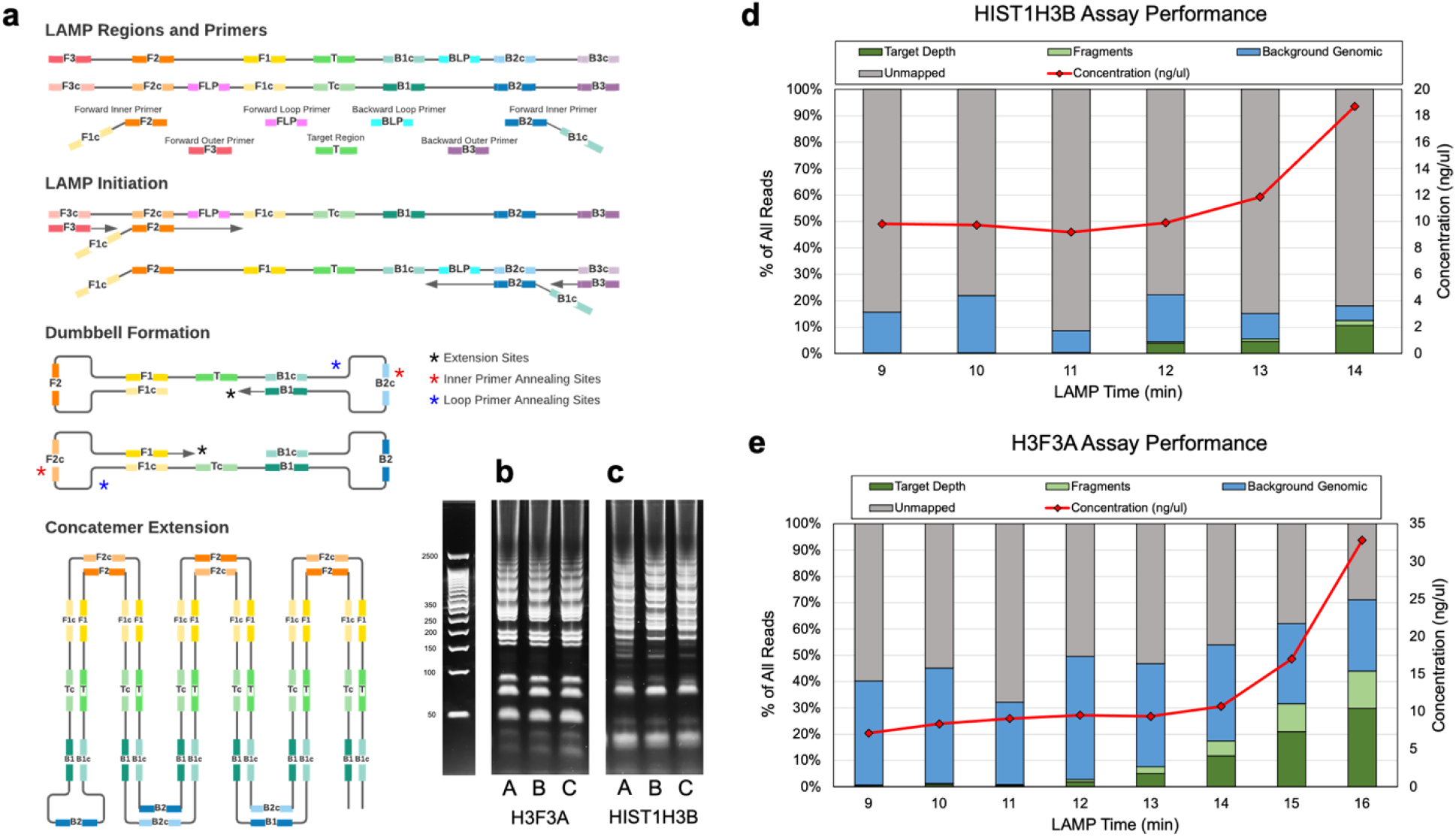
LAMP overview, and sequencing results from LAMP amplification timepoints. **(a)** LAMP amplifies target region of DNA and forms concatemers by leveraging a strand-displacing polymerase and primers that form intentional hairpin loops. Multiple different concatemer types can be formed and extended. **(b**,**c)** Gel-electrophoresis of six runs of LAMP product reveals many different types of product and potential primer dimers and primer sequences. **(b)** multiple runs on various input DNA samples shows the *H3F3A* assay forms clean bands, with concatemer types grouped according to integer multiples of the concatemer. The *HIST1H3B* product **(c)** displays similar patterns but seems contaminated by other spurious product and less organized. **(d)** DNA concentration measurements confirmed large amounts of amplification of the *HIST1H3B* assay at 14 minutes, but sequencing confirmed a large proportion of that product was not able to be mapped to the human genome, suggesting spurious amplification. **(e)** The *H3F3A* assay produced a larger proportion of target product indicating better assay behavior in-vitro.

To evaluate the feasibility of using LAMP as a replacement for PCR-based sequencing assays, we designed LAMP primer sets encompassing *H3F3A K27M* and *HIST1H3B K27M* hotspot mutations (see Methods section) and evaluated each for amplification rate and specificity. LAMP primer sets were designed such that the target hotspot mutation was maintained between the F2 and B2 regions (B1/B2 for the *H3F3A K27M* assay and F1/B1 for the *HIST1H3B K27M* assay) and not covered by the inner, or loop primers.

Approximate amplification rate was evaluated by starting multiple LAMP reactions at the same time and removing and quenching reactions in ice at 1-minute intervals (see Methods section). Specificity was difficult to evaluate using traditional gel electrophoresis due to the multiple possible types of LAMP base product and its concatemeric nature. However, proper LAMP concatemer formation should form clear groups of bands corresponding to integer multiple concatemer amplicons. Our *H3F3A* assay formed these clear groups, indicating proper amplification (Figure 5b) while our *HIST1H3B* assay formed fewer clear groups, indicating some spurious amplification or other partial assay malfunction (Figure 5c).

To measure LAMP product specificity more accurately, and identify an appropriate sequencing threshold time, we sequenced successive time-points of LAMP amplified product using various ONT barcodes from the ONT rapid barcoding library preparation kit (see Methods Section). Both assays induced amplification—increasing DNA concentration over time—however bioinformatic analysis (using the same method for PCR amplicon alignment and variant calling) revealed that much of the *HIST1H3B* product mass generated was unmappable (Figure 5d). In contrast, the *H3F3A* assay reliably amplified the target region and produced a more reasonable target fraction within 15 minutes (Figure 5e).

To further explore issues whether unmapped sequences were a failure of the LAMP assay, incompatibility between the LAMP product and ONT technology, or the applied bioinformatics toolchain, we designed a LAMP specific alignment algorithm—LAMPrey—to 1) diagnose the source of unmapped reads and 2) to recover useful target reads that standard long-read informatics pipelines are unable to align.

### LAMPrey: a Tool for LAMP Concatemer Analysis

LAMPrey relies on two main intuitions to accurately classify reads and properly identify diagnostically relevant target information: 1) properly formed LAMP concatemers will contain the LAMP sequences in their expected order, while 2) spurious amplification will contain many of these sequences, but in an improper order, and lacking target regions. LAMPrey begins by separately aligning the target region of interest, and all LAMP sequences (F3, F2, F1, B1, B2, B3) and their complements, to the read. All sequences that align to the read with at least 75% identity are marked as hits (Figure 6a). This step is analogous to “seeding” in common genomic read mappers; but here, LAMPrey is identifying seeds in the read from a small set of known, expected possibilities. If a target sequence is identified, LAMPrey looks to the left and right of the target for expected order of primer sequences according to proper LAMP amplification. Correct sequences are incorporated into a sub-read candidate for each identified target (Figure 6b). This step is analogous to seed “chaining” used by many read mapping algorithms. Once one or more sub-reads are identified, they are extracted and individually aligned to the region of interest using Minimap2^38^ (Figure 6c). If a sub-read aligns, it is assumed to be properly formed and can be used for variant calling. If more than one sub-read aligns to the target, a pileup is generated and considered as a group for variant calling (Figure 6d). A basecall is chosen via plurality of the calls in the pileup over the target. If there is a tie, a final call is randomly chosen between the tied plurality choices.

**Figure 6.**
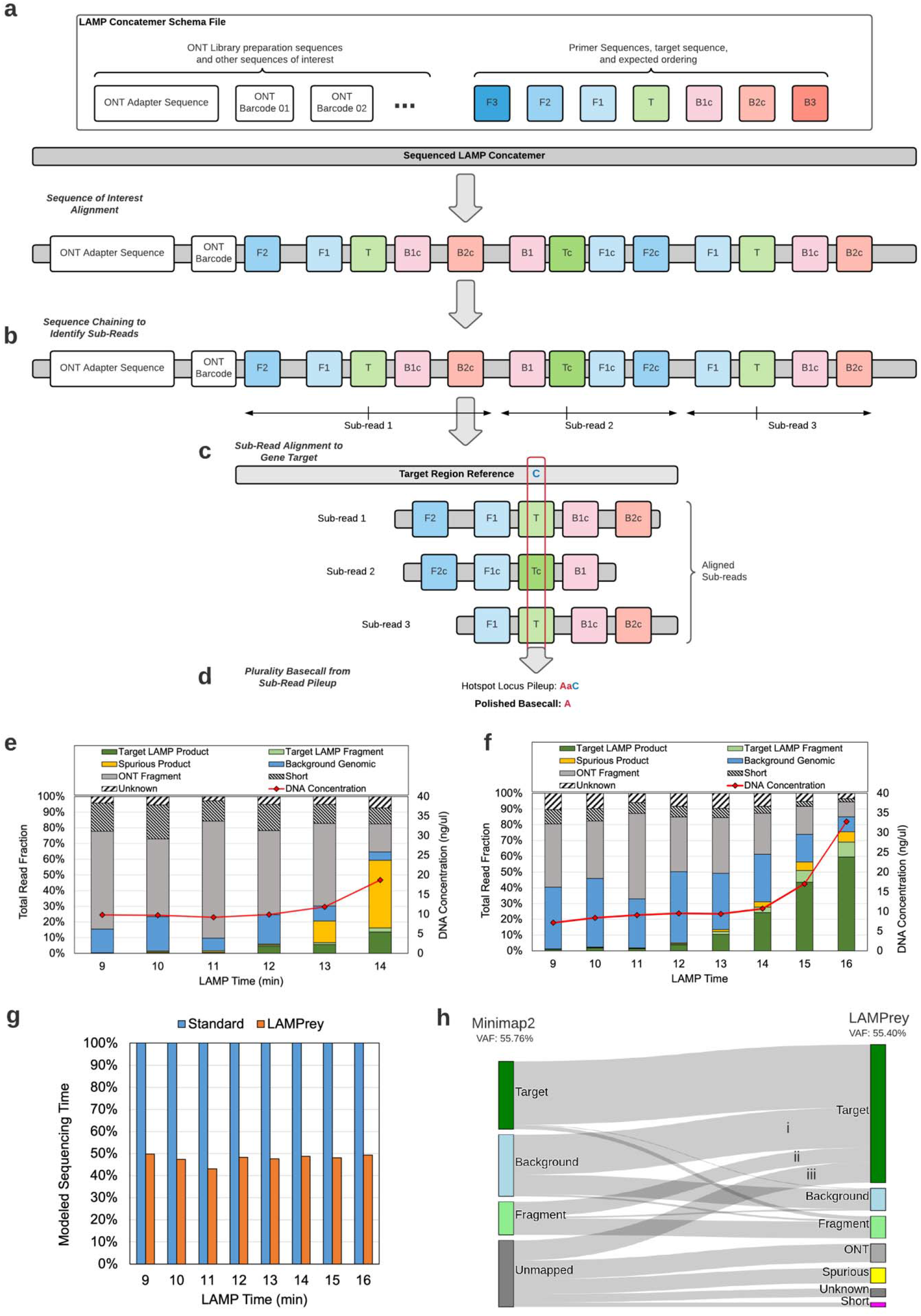
LAMPrey algorithm overview, classification results, and performance improvement relative to standard bioinformatics pipelines. **(a)** LAMPrey first marks suspected target regions and other sequences of interest in the candidate read. **(b)** Given a target instance, expected primer sequences are identified to the 5’ and 3’ ends of the target to form suspected concatemer sub-reads. **c** sub-reads are aligned to the gene target reference and **(d)** a consensus agreement is applied to define a call over the locus of interest. **(e**,**f)** LAMPrey is able to help diagnose spurious LAMP amplification or a properly functioning assay, and also **(g)** recover more diagnostically relevant reads than standard bioinformatics pipelines, leading to a reduction in sequencing time required to reach desired target variant call support. **(h)** Classifications of reads from the 16-minute *H3F3A* time-point using our standard pipeline (Minimap2) and LAMPrey. LAMPrey is able to recover a large amount of missed diagnostic information (i-iii) from reads missed by our standard pipeline. LAMPrey’s VAF (55.40%) is statistically identical to our standard approach (55.76%) indicating recovered information is not from an erroneous source.

If an alignable target sequence is identified, the read is classified as “Target.” If the read aligns to the LAMP region, but does not contain the target, it is classified as “Fragment.” If no target sequence is identified, but many primer sequences are found, we assume the LAMP process malfunctioned generating spurious product, and the read is classified as “Spurious”. If few or no potential primer sequences are identified in the read, and the read successfully maps to the human genome, it is classified as “Background” genomic DNA. Lamprey also identifies sequences which primarily contain ONT-related sequences such as adapter sequences and barcodes (classified as “ONT”) and short fragments (<60bp, classified as “Short”) that are difficult to classify. The remaining reads are classified as “Unknown.” A more detailed description of the LAMPrey algorithm is provided in Supplementary Materials.

We used LAMPrey to characterize sequenced LAMP product from 7,500 reads from each time point from the amplification time sweep runs of both *HIST1H3B* and *H3F3A* primer sets. For the *HIST1H3B* assay, LAMPrey was able to identify large amounts of spurious amplification (Figure 6e) which indicates a highly dysfunctional LAMP assay. In contrast, the *H3F3A* assay generates a consistently healthy proportion of on-target LAMP product and was chosen for final threshold sequencing evaluation (Figure 6f). However, it is worth noting that the *HIST1H3B* assay is still usable within the intra-operative timeframe, albeit with a time penalty due to the time wasted sequencing spurious product.

During assay diagnostics, we discovered that the LAMPrey tool was able to recover and align reads that otherwise would have been considered fragments, mapped incorrectly, or ended up unmapped when using our standard read alignment pipeline (see Methods section). When applied to the time-sweep sequencing results, LAMPrey was able to recover more than twice as many diagnostically relevant reads resulting in ∼2x improvement in modeled sequencing time-to-result (Figure 6g). To identify the source of benefit, we manually analyzed reads with diagnostically relevant information that Minimap2 missed, but LAMPrey was able to recover (Figure 6h, i-iii). This investigation revealed four major failure modes of our original informatics pipeline explaining LAMPrey’s improved performance:

1. *Off-target alignment, and filtering of non-primary alignments* (Figure 6e i): sequenced lamp concatemers are sometimes mapped to off-target locations. Correct alignments can be included as secondary or supplementary alignments by the mapper but are filtered out by the standard diagnostic pipeline.
2. *Imbalanced, fragmented concatemers* (Figure 6e ii): the fragmentation-based library preparation approach can leave concatemers imbalanced, where two different-sized sub-reads exist: one shorter with the target information, and one longer without. Read mapping algorithms will typically score the longer sub-read as the primary alignment, and soft clip the shorter, diagnostically relevant section.
3. *Short, hairpin reads* (Figure 6e iii): Concatemer fragments can contain short forward/reverse segments spanning the hairpin loops of a properly formed concatemer. The read mapper struggles to find proper chains for either of these short, competing sections, leading to mapping failure.
4. *Partially spurious, malformed concatemers* (Figure 6e iii): mis-primed LAMP amplification can occur after proper concatemer extension. If fragmentation generates a read that is mostly spurious, the read mapper may fail to align the read at all, even if a small section of target diagnostic information exists in the read.

LAMPrey did miss a small percentage of targets identified by our standard pipeline (Figure 6e). Upon further inspection, these reads were missed due to LAMPrey failing to identify target seeds in the initial seeding stage due to many basecalling errors. Tuning long-read aligner parameters and investigating one of many other alignment algorithms to improve performance is one possible approach to improve recovery of diagnostically relevant information. However, the complexity of LAMP concatemers coupled with a fragmentation-based library preparation approach and the above results clearly motivates a LAMP-specific analysis and further development of tools such as LAMPrey.

### LAMP-based Threshold Sequencing

We evaluated the feasibility of LAMP-based threshold sequencing by first conducting sequencing of various amplification times (Figure 7a). These libraries and the end-to-end sequencing demonstration were prepared using the optimized, 3-minute rapid DNA extraction protocol and 2-minute rapid adapter incubation library preparation protocol as previously identified as appropriate. Time-sweep sequencing and the Threshold Sequencing model identified 13-14 minutes as the time-optimal amplification target. LAMP amplification is so vigorous that small variations in initial conditions can lead to catastrophic under-amplification around the “optimal” amplification threshold, and unacceptably long sequencing times (Figure 7b). To account for this variability, we chose to evaluate a relatively conservative LAMP amplification time (14 minutes) to ensure a more predictable assay.

**Figure 7.**
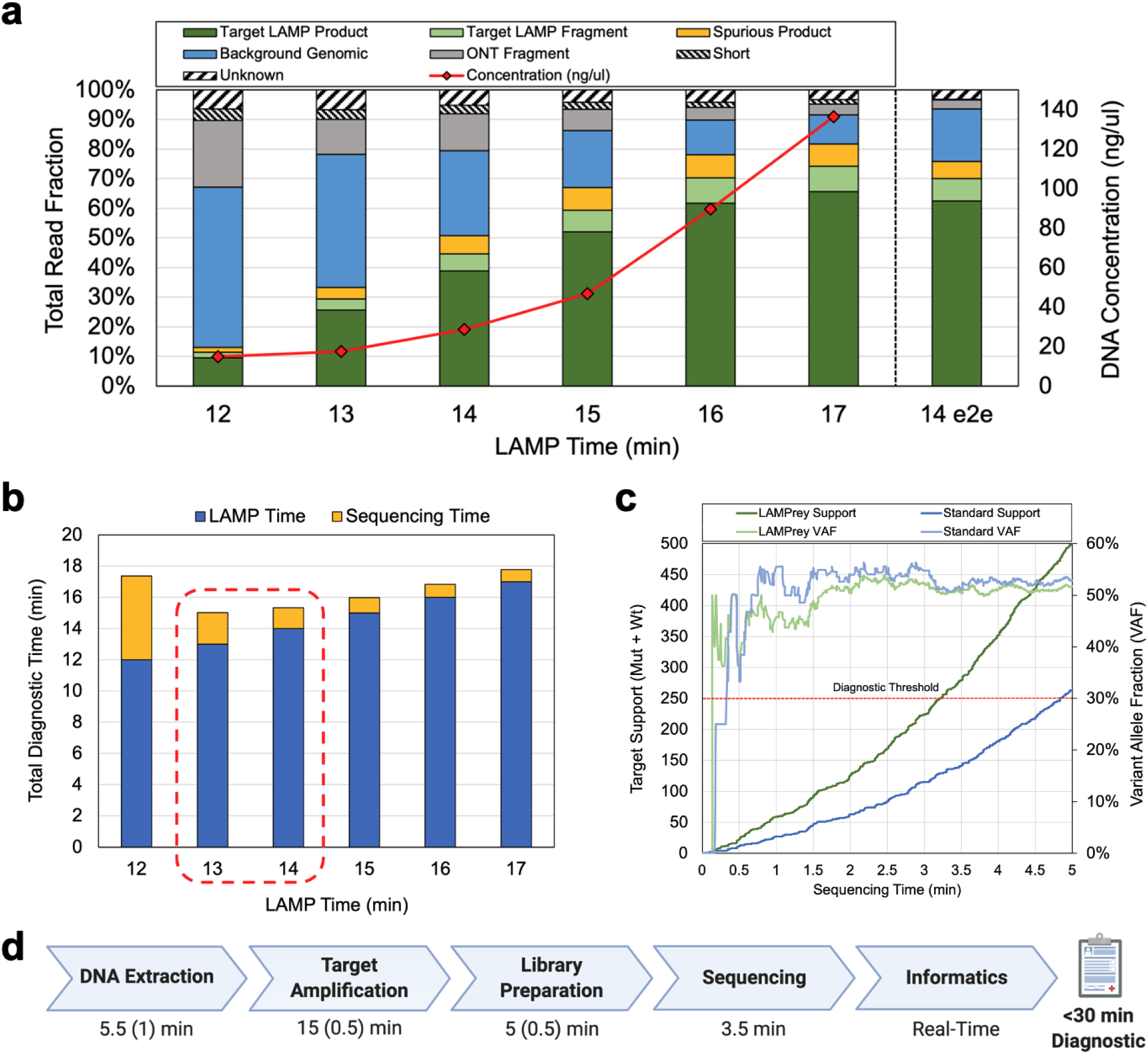
LAMP Threshold Sequencing assay results. **(a)** LAMPrey classification results from various amplification times and the final end-to-end demonstration (14 e2e) using DNA extracted from fresh tissue and the shortened LQE protocol. **(b)** Total sequencing time estimated from amplicon fraction at various LAMP time points. Optimal sequencing time is predicted to occur after 13-14 minutes of amplification. We chose to evaluate 14 minutes to guard against under-amplification. **(c)** Diagnostic results of the 14-minute end-to-end LAMP assay over time. A standard bioinformatics pipeline failed to leverage ∼48% of target reads. LAMPrey can recover this information and leads to a diagnosis ∼96 seconds sooner. **(d)** Approximate times for each diagnostic step with approximate manual overhead in parenthesis (Created with BioRender.com). Library preparation is notably faster than the 3-step PCR evaluation due to less hands-on-time, the introduction of ice quenching, and adapter incubation time optimization. Amplification time is lower due to our 14-minute LAMP protocol vs ∼26-minute (28 cycle) PCR amplification protocol.

We performed a timed, end-to-end run of the predicted time-optimal Threshold Sequencing protocol starting from an aliquot of ∼20mg of room-temperature tumor tissue, flash frozen at autopsy from a patient with previously diagnosed *H3F3A K27M* mutation. Diagnostically useful depth over time and sample variant allele fraction is shown in Figure 7c. Our protocol (see Methods section) was able to achieve 250x coverage and successfully call a known *H3F3A K27M* variant from a pediatric DIPG tumor sample in ∼29.5 minutes. The detailed protocol is shown in Supplementary Material (Figure S8) and available online (see Methods Section). Even with a well-behaved LAMP assay, a standard long-read alignment pipeline was not able to align some LAMP product reads that contained useful target information. Using the LAMPrey tool, we were able to recover ∼40% more information leading to a savings of ∼1.5 minutes of sequencing time (Figure 7c).

Amplification occurred more rapidly than expected, mimicking amplification expected at 16 minutes (Figure 7a). Model estimates again underestimated sequencing time required to reach 250x coverage (even with unexpectedly rapid amplification) due to a high proportion of available flow cell channels/pores not participating in sequencing during the first few minutes. Sequencing pores and pore sequencing rate was computed from the metadata in the Guppy basecalled fastq files (Methods). Only 264 pores participated in sequencing within the first 3.5 minutes, even though ∼430 pores were reported healthy and available for sequencing. Each active channel sampled reads at a rate almost identical to our model’s value (0.0165 reads/channel/s vs 0.0158 reads/channel/s) indicating that each pore’s access to sequence-able strands immediately after loading was a limiting factor in proper prediction and achieving a more rapid time-to-result.

The approximate breakdown of times for each step, including human overhead, is shown in Figure 7d. The detailed protocol is shown in Supplementary Material (Figure S8) and available online (see Methods Section). To the best of our knowledge, this is the first sub-30-minute sequencing-based variant call achieved to date and demonstrates the feasibility of using sequencing as a diagnostic technique within the intra-operative timeframe.

## Discussion

Threshold Sequencing provides the first demonstration of the feasibility of sequencing-based molecular diagnostics within the intraoperative timeframe (<1hr). By co-optimizing DNA extraction, target amplification, library preparation, and informatics, we were able to achieve a clinically-relevant, sequencing-based variant call from real patient tissue within 30 minutes.

By using the LAMPrey tool to polish LAMP concatemers with multiple align-able target regions, we were able to filter out low-confidence reads and calls. This self-polishing feature could be leveraged in the future to provide highly accurate variant calls at lower coverage, and for use-cases where variant allele fractions are near or lower than the ONT sequencer error rate (e.g. liquid biopsy).

While this work focused on designing assays for *HIST1H3B K27M* and *H3F3A K27M* histone mutations—common in pediatric diffuse midline gliomas—the Threshold Sequencing technique can be applied to any assay and target other hotspot mutations or oncogenes. Mutation hotspots such as *BRAF V600, IDH1 R132*, and *IDH2 R172*, and *IDH2 R140* are prime examples where identifying mutational status during surgery could inform more-aggressive resection (*BRAF*) or less-aggressive resection (*IDH1/2*). Intra-operative sequencing could help augment current intra-operative diagnostics such as frozen section histology, by providing quantitative support for a qualitative—and perhaps indeterminant—histological evaluation.

The Threshold Sequencing protocols are low-complexity and all required equipment could easily fit onto a surgical cart. This would allow the assay to be practically performed within an operating room or adjoining pathological suite without the overhead of moving the sample to a dedicated diagnostic laboratory. We also expect these protocols to be automatable and performable using laboratory robots or microfluidic labs-on-a-chip (e.g. ONT’s Voltrax device). Automation would further reduce expertise required to perform the assay, reduce human error and inefficiencies, and reduce total required equipment footprint.

Threshold sequencing currently relies on pre-characterizing amplification rates to decide on a proper amplification threshold for a particular tissue type. While we expect this pre-characterization approach to work well over multiple patients and samples, future implementations could leverage fluorescence or other real-time amplification measurement to identify the optimal sequencing threshold cutoff point (based on a pre-characterized expected *F*_*target*_) in real-time. This would greatly enhance the reliability and reproducibility of the assay, given various input types and/or other variables that could impact amplification reliability.

While our current realization is a singleplex assay and requires 14-minutes of amplification time, higher throughput devices on ONTs future product roadmap—with orders of magnitude more parallel pores—would allow for much lower sequencing thresholds and time-to-result, and/or highly multiplexed intra-operative assays. With the use of barcodes, multiple tumor locations could be sequenced intra-operatively to provide a matched tumor/normal sample for more accurate variant calling, copy number variation analysis, or even a spatial map of tumor genetics, quantitatively outlining tumor margin and mapping potential molecular heterogeneity intraoperatively.

## Acknowledgements

This work was funded in part by the US National Science Foundation (NSF) grant NSF 1652294, and NSF 2030454, and the D. Dan and Betty Kahn Foundation.

## Methods

### PCR primer design and size evaluation

PCR primer sets were drawn from multiple sources and selected or designed to flank the *HIST1H3B K27M* hotspot mutation. The 187 bp primer set was taken from prior work^39^ . The 260 bp primer set was designed using Primer3 v4.1.0 and tailed at the 5’ ends with ONT handshake sequences to enable potential interoperability with ONT’s four-primer library preparation protocol. The 609bp and 910bp primer sets were designed by Integrated DNA Technologies. All primers were supplied by Integrated DNA technologies. PCR was performed using New England Biolabs (NEB) Q5 2x Master mix (NEB #M0492) and standard primer concentrations. 30ng of NA12878 DNA (Coriell Institute) was used as template and 35 cycles of PCR was performed on a Biorad C1000 thermocycler with the following cycling parameters: Initial denaturation for 30s @ 98°C, 35 cycles of 10s @ 98°C, 15s @ 58°C-68°C, 40s @ 72°C, and a final extension at 72°C for 2 minutes. Library preparation was performed using Oxford Nanopore’s Rapid Barcoding Kit (ONT #SQK-RBK004) and sequenced using a MinION R9.4.1 flow cell. Fast5 signal files were basecalled using ONT’s GPU accelerated basecaller “Guppy” version 4.2.2. Reads were aligned using Minimap2 version 2.17 (-x map-ont) against the human reference genome (GRCh37/hg19). Alignments were filtered for primary alignments using samtools^40^ v1.7 (-F 0×900). Alignments were classified using according to a custom script (see Code Availability). All primer sets are listed in Supplementary Materials.

### DNA Extraction

DNA was extracted from aliquots of two pediatric DIPG patients (one *HIST1H3B K27M* positive, one *H3F3A K27M* positive) originally acquired at autopsy, flash frozen, and stored at -80°C. Sub-aliquots were divided into sections between 20-50mg and stored at -20°C before experimentation. DNA extraction was performed after tissue was allowed to thaw to room temperature. For PCR experiments targeting the *H3IST1H3B K27M* variant, we used Lucigen QuickExtract DNA extraction solution (Lucigen #QE0905T) and the standard protocol was followed. ∼20-30mg of tissue was placed in 500ul of Lucigen QuickExtract solution in a 2ml Eppendorf tube and vortexed for 15s. The solution was incubated for 6 minutes at 65°C, vortexing briefly after 3 minutes and 6 minutes. Solution was finally incubated at 98°C for 2 minutes before either being 1) immediately used for amplification or 2) stored at 4°C for later amplification. For LAMP experiments targeting the *H3F3A K27M* variant, ∼20-30mg of tissue was placed in 200ul Lucigen QuickExtract solution in a 1.5ml tube and processed as above.

The Lucigen QuickExtract Protocol was optimized by shrinking the incubation time to 1 minute and increasing the initial vortex time to 30s and eliminating intermittent vortexing steps. The optimized protocol also uses 100ul of LQE per ∼20mg aliquot of tissue and uses 0.2ul PCR tubes. This allows the user to use one thermocycler for extraction, amplification, and library preparation, obviating the use of multiple heat blocks. This optimized methodology was used in the end-to-end LAMP experiment.

### PCR cycle-sweep amplification and sequencing

PCR reactions were prepared according to the NEB Q5 2x master mix (NEB #M0492) protocol using a 1ul of LQE extracted DNA product as input (∼20ng DNA). PCR of varying cycles was run sequentially on a BioRad C1000 thermocycler with a 3 °C/s ramp rate with the following cycling parameters for 20, 22, 24, 26, 28, and 30 cycles: Initial denaturation for 30s @ 98°C, *N* cycles of 5s @ 98°C, 5s @ 64°C, 8s @ 72°C, and a final extension at 72°C for 2 minutes. The sequencing library was prepared according to the ONT rapid barcoding kit protocol (ONT #SQK-RBK004) with separate barcodes assigned for each PCR cycle number and sequenced on a MinION R9.4.1 flow-cell. Resulting Fast5 signal files were basecalled using ONT’s GPU accelerated basecaller Guppy version 4.2.2. Reads were aligned using Minimap2 version 2.17 (-x map-ont) against the human reference genome (GRCh37/hg19). Alignments were filtered for primary alignments using samtools v1.7 (-F 0×900) and classified according to alignment position and pileup results using a custom script (see Code Availability). Post-amplification DNA concentration was measured via Qubit (Invitrogen dsDNA HS assay #Q33230).

### PCR End-to-end Threshold Sequencing

A full protocol is available at https://www.protocols.io/view/ultra-rapid-sequencing-pcr-bs7bnhin. Briefly, tumor tissue was allowed to thaw and come to room temperature. DNA extraction was performed (see Methods Section) with the run timer started when the LQE liquid first comes into contact with the tissue. After extraction and a 30 second ice quench, 1ul of extracted DNA was added to a tube with 24ul pre-mixed PCR master mix, primers, and water (according to the standard 25ul Q5 2x Master Mix protocol) and placed in a thermocycler for amplification. During amplification, a sequencing run in the ONT MinKNOW software was started and the MinION flow cell was initiated (pore health check and initial mux scan) and primed. A sequencing run was started without a library and paused using the MinKNOW software. After the PCR protocol finished, 7.5ul of PCR product was immediately added to 2.5ul of fragmentation mix and tagmentation was initiated in the same thermocycler according the ONT Rapid Library Preparation protocol (ONT #SQK-RAD004). Tagmented product was allowed to be brought down to room temperature by the thermocycler. Rapid sequencing adapter was added (1ul), gently pipetted to mix, and allowed to incubate for 5 minutes. During this time, the ONT sequencing mix was prepared according to the standard protocol. With 1-minute remaining, the flow cell was re-primed for library loading. After 5-minute incubation, the library was then immediately mixed with the sequencing mix via pipetting to ensure loading bead/library contact and loaded onto the flow-cell. The run was then immediately re-started in the MinKNOW software to initiate sequencing. Informatics (basecalling, alignment, variant calling) was performed after the fact but verified to be performable in real-time. Resulting reads were basecalled, aligned, and filtered using the standard approach. Time to variant call was computed from FASTQ reads sorted by ONT read start time timestamps and extracted using a custom script (see Code Availability). For this off-line analysis, we assume the start time approximates data availability given that a vast majority of sequenced reads take <1 second to sequence, and the rapidity of basecalling/analysis.

### LAMP primer design and time sweep

LAMP primer sets were designed using Premier Biosoft LAMP Designer v1.16 and evaluated for efficiency and specificity using both gel electrophoresis and sequencing. Care was taken to pick primer sets where the hotspot mutation of interest (*H3K27M*) was not covered by primers and present between the F2 and B2 LAMP regions. LAMP amplification was performed using NEB’s WarmStart 2x LAMP kit (NEB #E1700) following the standard protocol but omitting the final polymerase deactivation before sequencing. To measure amplification over time, separate reactions were prepared using the *HIST1H3B* primer set and run simultaneously. Reactions were pulled at various time points minutes and immediately quenched in ice water for 15s to stop amplification. The sequencing library was prepared according to the ONT rapid barcoding kit protocol (ONT #SQK-RBK004) assigning separate barcodes to each time-point and sequenced on a MinION R9.4.1 flow-cell. Resulting Fast5 signal files were basecalled using ONT’s GPU accelerated basecaller Guppy version 4.2.2. Reads were aligned using Minimap2 version 2.17 (-x map-ont) against the human reference genome (GRCh37/hg19). Alignments were filtered for primary alignments using samtools v1.7 (-F 0×900). Post-amplification DNA concentration was measured via Qubit (Invitrogen dsDNA HS assay #Q33230).

### LAMP Threshold Sequencing

Prior to beginning, the MinION flow cell was initiated (pore health check and initial mux scan) and primed and LAMP, tagmentation mix, and sequencing buffers were all prepared. A sequencing run was started without a library and paused using the MinKNOW software. 1ul of DNA (∼29ng) from patient tumor tissue extracted according to the standard Lucigen QuickExtract DNA Extraction protocol was sampled at room temperature and added to a tube with pre-mixed NEB WarmStart LAMP 2x Master Mix (NEB #E1700), *H3F3A* primer set, and nuclease free water according to the NEB protocol. LAMP was performed in a thermocycler at 65C for 14 minutes, followed by a 15s quench in ice water. After amplification, 1.9ul LAMP product was added to 2.5ul of ONT fragmentation mix (FRA) and 5.6ul of nuclease free water and tagmentation was initiated in the same thermocycler according to the ONT protocol (ONT #SQK-RAD004). Tagmented product was quenched in ice water for 15s to bring to room temperature. 1ul of ONT rapid sequencing adapter (RAP) was added, gently pipetted to mix, and allowed to incubate for 2 minutes. The DNA library was then immediately mixed with the sequencing mix via pipetting and loaded onto the flow-cell according to the standard protocol. The flow-cell was then immediately re-started to initiate sequencing. Informatics (basecalling, alignment, variant calling) was performed after the fact but verified to be performable in real-time. Resulting Fast5 signal files were basecalled using ONT’s GPU accelerated basecaller Guppy version 4.2.2. Reads were aligned using Minimap2 version 2.17 (-x map-ont) against the human reference genome (GRCh37/hg19) or processed using the LAMPrey tool. A full detailed protocol is available at https://www.protocols.io/view/ultra-rapid-sequencing-lamp-btvmnn46. Available/active pores and per channel sequencing rate was computed using the sequencing rate script (see Code Availability). Informatics (basecalling, alignment, variant calling) was performed after the fact but verified to be performable in a pipelined fashion in real-time. Resulting reads were basecalled, aligned, and filtered using the standard approach. Time to variant call was computed from FASTQ reads sorted by ONT read start time timestamps and extracted using a custom script or via the LAMPrey tool (see Code Availability). For this off-line analysis, we assume the Guppy basecaller *start_time* approximates data availability and variant call given that a vast majority of sequenced reads take <1 second to sequence, and the rapidity of basecalling/analysis.

## Data Availability

All sequencing data (fastq) generated from the timed end-to-end runs has been uploaded to the NCBI SRA database under accessions SAMN19108851 (PCR) and SAMN19108852 (LAMP). Fast5 files and other data are available on request.

## Code Availability

All scripts and code used in this evaluation are available in the following GitHub repository: https://www.github.com/jackwadden/UltraRapidSeq. The LAMPrey tool is available in the following GitHub repository: https://www.github.com/jackwadden/lamprey.

## References

1. Louis, D. N. et al. The 2016 World Health Organization Classification of Tumors of the Central Nervous System: a summary. Acta Neuropathol. (Berl.) 131, 803–820 (2016).

2. Histone H3K27M Mutation Overrides Histological Grading in Pediatric Gliomas | Scientific Reports. https://www.nature.com/articles/s41598-020-65272-x.

3. Lu, V. M., Alvi, M. A., McDonald, K. L. & Daniels, D. J. Impact of the H3K27M mutation on survival in pediatric high-grade glioma: a systematic review and meta-analysis. J. Neurosurg. Pediatr. 23, 308–316 (2018).

4. Wang, L. et al. PIK3CA mutations frequently coexist with EGFR/KRAS mutations in non-small cell lung cancer and suggest poor prognosis in EGFR/KRAS wildtype subgroup. PloS One 9, e88291 (2014).

5. Gojo, J. et al. Personalized Treatment of H3K27M-Mutant Pediatric Diffuse Gliomas Provides Improved Therapeutic Opportunities. Front. Oncol. 9, (2020).

6. Chen, R. et al. Molecular features assisting in diagnosis, surgery, and treatment decision making in low-grade gliomas. Neurosurg. Focus 38, E2 (2015).

7. Toomey, S. et al. Identification and clinical impact of potentially actionable somatic oncogenic mutations in solid tumor samples. J. Transl. Med. 18, 99 (2020).

8. Kanamori, M. et al. Rapid and sensitive intraoperative detection of mutations in the isocitrate dehydrogenase 1 and 2 genes during surgery for glioma. J. Neurosurg. 120, 1288–1297 (2014).

9. Koriyama, S. et al. A surgical strategy for lower grade gliomas using intraoperative molecular diagnosis. Brain Tumor Pathol. 35, 159–167 (2018).

10. Liu, Z. et al. Intraoperative Chemotherapy with a Novel Regimen Improved the Therapeutic Outcomes of Colorectal Cancer. J. Cancer 10, 5986–5991 (2019).

11. Terata, K. et al. Novel rapid-immunohistochemistry using an alternating current electric field for intraoperative diagnosis of sentinel lymph nodes in breast cancer. Sci. Rep. 7, 2810 (2017).

12. Huang, T. et al. Detection of histone H3 K27M mutation and post-translational modifications in pediatric diffuse midline glioma via tissue immunohistochemistry informs diagnosis and clinical outcomes. Oncotarget 9, 37112–37124 (2018).

13. Livermore, L. J. et al. Rapid intraoperative molecular genetic classification of gliomas using Raman spectroscopy. Neuro-Oncol. Adv. 1, vdz008 (2019).

14. Chrzanowska, N. M., Kowalewski, J. & Lewandowska, M. A. Use of Fluorescence In Situ Hybridization (FISH) in Diagnosis and Tailored Therapies in Solid Tumors. Molecules 25, (2020).

15. Duncan, D. J., Vandenberghe, M. E., Scott, M. L. J. & Barker, C. Fast fluorescence in situ hybridisation for the enhanced detection of MET in non-small cell lung cancer. PLOS ONE 14, e0223926 (2019).

16. Al-Ramadhani, S. et al. Metasin—An Intra-Operative RT-qPCR Assay to Detect Metastatic Breast Cancer in Sentinel Lymph Nodes. Int. J. Mol. Sci. 14, 12931–12952 (2013).

17. Ferris, R. L. et al. Intraoperative qRT-PCR for Detection of Lymph Node Metastasis in Head and Neck Cancer. Clin. Cancer Res. Off. J. Am. Assoc. Cancer Res. 17, 1858–1866 (2011).

18. Smith, J. S. et al. Alterations of Chromosome Arms 1p and 19q as Predictors of Survival in Oligodendrogliomas, Astrocytomas, and Mixed Oligoastrocytomas. J. Clin. Oncol. 18, 636–636 (2000).

19. Magi, A. et al. Nano-GLADIATOR: real-time detection of copy number alterations from nanopore sequencing data. Bioinforma. Oxf. Engl. 35, 4213–4221 (2019).

20. High resolution copy number inference in cancer using short-molecule nanopore sequencing | bioRxiv. https://www.biorxiv.org/content/10.1101/2020.12.28.424602v1.

21. Sallman, D. A. & Padron, E. Integrating mutation variant allele frequency into clinical practice in myeloid malignancies. Hematol. Oncol. Stem Cell Ther. 9, 89–95 (2016).

22. Berger, M. F. & Mardis, E. R. The emerging clinical relevance of genomics in cancer medicine. Nat. Rev. Clin. Oncol. 15, 353–365 (2018).

23. Euskirchen, P. et al. Same-day genomic and epigenomic diagnosis of brain tumors using real-time nanopore sequencing. Acta Neuropathol. (Berl.) 134, 691–703 (2017).

24. Notomi, T. et al. Loop-mediated isothermal amplification of DNA. Nucleic Acids Res. 28, e63 (2000).

25. Kashir, J. & Yaqinuddin, A. Loop mediated isothermal amplification (LAMP) assays as a rapid diagnostic for COVID-19. Med. Hypotheses 141, 109786 (2020).

26. LamPORE: rapid, accurate and highly scalable molecular screening for SARS-CoV-2 infection, based on nanopore sequencing | medRxiv. https://www.medrxiv.org/content/10.1101/2020.08.07.20161737v3.

27. Chou, P.-H., Lin, Y.-C., Teng, P.-H., Chen, C.-L. & Lee, P.-Y. Real-time target-specific detection of loop-mediated isothermal amplification for white spot syndrome virus using fluorescence energy transfer-based probes. J. Virol. Methods 173, 67–74 (2011).

28. Guo, X.-G. et al. Evaluation of the real-time fluorescence loop-mediated isothermal amplification assay for the detection of Streptococcus agalactiae. Biosci. Rep. 39, (2019).

29. Gill, P. & Hadian Amree, A. AS-LAMP: A New and Alternative Method for Genotyping. Avicenna J. Med. Biotechnol. 12, 2–8 (2020).

30. Hardinge, P. & Murray, J. A. H. Reduced False Positives and Improved Reporting of Loop-Mediated Isothermal Amplification using Quenched Fluorescent Primers. Sci. Rep. 9, 7400 (2019).

31. AmpliSeq for Illumina Focus Panel | Combined DNA and RNA workflow. https://www.illumina.com/products/by-type/sequencing-kits/library-prep-kits/ampliseq-focus-panel.html.

32. Rapid Sequencing Kit (RAD004). https://store.nanoporetech.com/us/sample-prep/rapid-sequencing-kit.html.

33. Picelli, S. et al. Tn5 transposase and tagmentation procedures for massively scaled sequencing projects. Genome Res. 24, 2033–2040 (2014).

34. Nextera XT Library Prep: Tips and Troubleshooting. 6.

35. DNeasy Blood & Tissue Kits - QIAGEN Online Shop. https://www.qiagen.com/us/products/top-sellers/dneasy-blood-and-tissue-kit/#orderinginformation.

36. QuickExtract™ DNA Extraction Solution | Lucigen. https://www.lucigen.com/QuickExtract-DNA-Extraction-Solution/.

37. Ding, S. et al. One-step colorimetric genotyping of single nucleotide polymorphism using probe-enhanced loop-mediated isothermal amplification (PE-LAMP). Theranostics 9, 3723–3731 (2019).

38. Minimap2: pairwise alignment for nucleotide sequences | Bioinformatics | Oxford Academic. https://academic.oup.com/bioinformatics/article/34/18/3094/4994778.

39. Bruzek, A. K. et al. Electronic DNA Analysis of CSF Cell-free Tumor DNA to Quantify Multi-gene Molecular Response in Pediatric High-grade Glioma. Clin. Cancer Res. 26, 6266–6276 (2020).

40. Sequence Alignment/Map format and SAMtools | Bioinformatics | Oxford Academic. https://academic.oup.com/bioinformatics/article/25/16/2078/204688.

