## Supplementary Material for "Ultra-Rapid Somatic Variant Detection via Real-Time Threshold Sequencing"

**PCR and LAMP Primer Sets, Melting Temperatures, and Cycling Parameters**

Primer sets used in all evaluations are listed in Table S1 below. ONT handshake sequence tails are highlighted in red and blue. For tailed primers, both the genomic length and total amplicon length including the handshake sequences (bolded) are listed. For FIP and BIP primers, poly-T bend segments are bolded.

| **Primer Set** | **Genome Coverage** | **Amplicon Size** | **Name** | **Sequence** |
| --- | --- | --- | --- | --- |
| 3-step 146 | 146 | 190 | Fwd | **TTTCTGTTGGTGCTGATATTGC**CTGCAGGCAAGCTTTTCT |
|  |  |  | Rev | **ACTTGCCTGTCGCTCTATCTTC**AGGCTTTTTCACGCCG |
| 3-step 216 | 216 | 260 | Fwd | **TTTCTGTTGGTGCTGATATTGC**GAATCAGCAACTCGGTCGAC |
|  |  |  | Rev | **ACTTGCCTGTCGCTCTATCTTC**CAGGCAAGCTTTTCTGTGGT |
| 3-step 565 | 565 | 609 | Fwd | **TTTCTGTTGGTGCTGATATTGC**AGCTTCCGAATCAGCAACTC |
|  |  |  | Rev | **ACTTGCCTGTCGCTCTATCTTC**TTCCCTAATCGTCTCCAATATTCTC |
| 3-step 910 | 866 | 910 | Fwd | **TTTCTGTTGGTGCTGATATTGC**CTCCTGCAGCGCCATCA |
|  |  |  | Rev | **ACTTGCCTGTCGCTCTATCTTC**AGTCTCCCTCTCGCCCAA |
| 2-step | 254 | 298 | Fwd | **TTTCTGTTGGTGCTGATATTGC**AGGCTTTTTCACGCCGCCGGTA |
|  |  |  | Rev | **ACTTGCCTGTCGCTCTATCTTC**GGACCAATCCAAGAAGGGCGCGGGG |
| Pig TP53 |  |  | Fwd | **TTTCTGTTGGTGCTGATATTGC**TCTGTTCGCTCTCCATCCTC |
|  |  |  | Rev | **ACTTGCCTGTCGCTCTATCTTC**CCACCTCGGTCATGTACTCT |
| HIST1H3B LAMP |  |  | F3 | GATCGGTCTTGAAGTCTTGG |
|  |  |  | B3 | ATAGTTGGTGGTCTGACTCTAT |
|  |  |  | FIP | AAAGCCTCACCGTTACCG**TTTT**CCGAATCAGCAACTCG |
|  |  |  | BIP | GAGCAGCCTTGGTAGCCAG**TTTT**GGCTCGTACTAAACAGACAG |
|  |  |  | FLP | GAGATCCGCCGCTACCAA |
|  |  |  | BLP | TTACCGCCGGTGGATTTC |
| H3F3A LAMP |  |  | F3 | GTTTGGTAGTTGCATATGGTG |
|  |  |  | B3 | ATACCTGTAACGATGAGGTTTC |
|  |  |  | FIP | GCGGGCAGTCTGCTTTGTA**TTTT**ATGCTGGTAGGTAAGTAAGGA |
|  |  |  | BIP | CGACCGGTGGTAAAGCACC**TTTT**CACCCCTCCAGTAGAG |
|  |  |  | FLP | CGAGCCATGGTACAGAGAC |
|  |  |  | BLP | CAGGAAGCAACTGGCTACA |

Table S1. **All primer sequences used in experiments in the paper and supplementary materials**.

| **Experiment** | **Initial Denature** | **Cycles** | **Denature** | **Anneal** | **Extend** | **Final Extend** |
| --- | --- | --- | --- | --- | --- | --- |
| 3-step 146 length eval | 30s @ 98°°C | 35 | 10s @ 98°C | 15s @ 67.8°C | 40s @ 72°C | 120s @ 72°C |
| 3-step 216 length eval | 30s @ 98°C | 35 | 10s @ 98°C | 15s @ 64°C | 40s @ 72°C | 120s @ 72°C |
| 3-step 565 length eval | 30s @ 98°C | 35 | 10s @ 98°C | 15s @ 65°C | 40s @ 72°C | 120s @ 72°C |
| 3-step 866 length eval | 30s @ 98°C | 35 | 10s @ 98°C | 15s @ 70.7°C | 40s @ 72°C | 120s @ 72°C |
| 2-step eval | 30s @ 98°C | 28 | 5s @ 98°C | 10s @ 72°C | - | 30s @ 72°C |
| 3-step 216 end-to-end | 30s @ 98°C | 28 | 5s @ 98°C | 5s @ 64°C | 8s @ 72°C | 30s @ 72°C |
| 4PP 3-step 216 eval | 30s @ 98°C | 35 | 5s @ 98°C | 10s @ 64°C | 13 @ 72°C | 30s @ 72°C |
| LAMP end-to-end | - | - | 14m @ 65°C | - | - | - |

Table S2. **Cycling parameters used for all experiments.**

**Survey of Prior Ultra-Rapid Molecular Diagnostics**

| **Technique** | **Detection Method** | **Detectable Cancer Markers** | **Reportable Target Mutations** | **Fastest End-to-end Result** | **Targeted Oligonucleotide**  **Amplification** | **Quantitative Variant Fraction**  **(*inferred)** | **Streaming Output** |
| --- | --- | --- | --- | --- | --- | --- | --- |
| IHC | Stain | Tumor Associated Antigen | 0 | 20-30 mins^1,2^ | n | n | n |
| FISH | Stain | Allele specific mutation, CNVs | - | 65 min^3^ | n | n | n |
| Raman Spectroscopy | Spectroscopy | *IDH* Mutant or not | - | 15 min^4^ | n | n | n |
| RT-qPCR | Fluorescence | Gene Expression | - | 32-42 min^5,6^ | y | n | y |
| qPCR | Fluorescence | Allele Specific Point Mutation, CNVs | 2-7 | 60-65 min^7,8^ | y | y* | y |
| Sanger | Sequencing | Point mutations, CNVs | **4^n | 4.5hr^7^ | y | y | n |
| Illumina | Sequencing | Point mutations, CNVs | **4^n | 17 hrs^9^ | y | y | n |
| PacBio SMRT | Sequencing | Point mutations, CNVs | **4^n | >4hrs | y | y | y |
| ONT MinION Ligation | Sequencing | Point mutations, CNVs | **4^n | 2.5hrs^10^ | y | y | y |
| **This Work** | **Sequencing** | Point mutations, **CNVs | ****4^n** | **<30 min** | **y** | **y** | **y** |

Table S3. **Molecular diagnostic techniques and associated rapid time-frames**. Sequencing-based diagnostics have not been accomplished within the intra-operative time-frame. *qPCR variant allele fractions are inferred from comparison to wildtype allele amplification or a standard curve and have not been demonstrated intra-operatively. **Sequencing can identify any mutation that lies within an amplicon; therefore, the number of reportable target mutations is at least 4^*n* (where ‘*n*’ is the length of the amplicon(s) under evaluation*).* ***This work does not evaluate copy number variation detection, but should be capable of detecting these types of mutations given the addition of an assay targeting a copy-number control gene. FISH = Fluorescent In-Situ Hybridization; IHC = Immunohisto Chemistry; RT-qPCR = reverse transcription quantitative polymerase chain reaction; qPCR = quantitative polymerase chain reaction.

**Threshold Sequencing Model Example Parameters**

Figure 3C develops a model framework to analytically derive the optimal amount of time spent on amplification in order to guarantee minimal total diagnostic time. The equations used to build the model and the corresponding input parameters used to generate Figure 3C are shown below in Table S4.

| **Parameter** | **Description** | **Value or Equation** |
| --- | --- | --- |
| $N_{depth}$ | Diagnostic coverage requirement | 250x |
| $T_{cycle}$ | PCR Cycle Time | 60s |
| $T_{init}$ | PCR initial denaturation time | 60s |
| $T_{final}$ | PCR final extension time | 60s |
| $N_{pores}$ | Active Channels/Pores Contributing to Sampling | 512 |
| $R_{seq}$ | Transposition rate of DNA through the Nanopore | 400bp/s |
| $T_{capture}$ | Average Capture Time (time-between read capture per pore) | 2s |
| $L_{background}$ | Average Background Read Length | 4kbp |
| $L_{target}$ | Target amplicon length | 250bp |
| $L_{background}$ | Background Genomic Size | 3,079,843,747bp |
| $N_{background}$ | The number of background reads | $N_{background}=\frac{S_{background}}{S_{read}}$ |
| $N_{target}$ | The number of targets | $N_{target}= 2^{cycles}$ |
| $F_{target}$ | The fraction of target amplicons relative to background genomic reads | $F_{target}=\frac{N_{target}}{N_{background}+ N_{target}}$ |
| $L_{read}$ | Average sequenced read length | $L_{read}= F_{target}* L_{target}+ \left( 1-F_{target} \right)*L_{background}$ |
| $R_{sample}$ | Average sampling rate (reads/pore/second) | $R_{sample}=1/(\frac{L_{read}}{R_{seq}}+T_{capture})$ |
| $T_{seq}$ | Time to sequence to target depth | $T_{seq}=N_{depth}*(\frac{1}{N_{pores}*R_{sample}*F_{target}})$ |
| $T_{amp}$ | Amplification time (PCR) | $T_{amp}=T_{init}+T_{cycle}*N_{cycle}+T_{final}$ |
| $T_{total}$ | Total diagnostic time (amplification + sequencing) | $T_{total}=T_{amp}+T_{seq}$ |

Table S4. **Model parameters and equations used to generate Figure 3C.**

**Impact of Rapid Adapter Incubation Time**

We tested four different rapid adapter incubation times from 5 minutes to 2 minutes. Five minutes is the incubation time recommended by ONT’s Rapid Barcoding library preparation kit (SQK-RBK004). Reactions were started at successive 1-minute intervals to assure all incubations ended at the same time. Once incubations finished, each reaction was immediately and independently mixed with ¼ ONT pre-mixed sequencing mix. This was to prevent ONT loading beads from preferentially binding to amplicons first added to one pooled sequencing mix, as observed in earlier experiments. All reactions were then mixed, combined, and sequenced until the lowest target depth reached >5,000 target reads. Fast5 files were basecalled using Guppy V4.2.2 and aligned using Minimap^11^ v2.17. Adapter efficiency was estimated by comparing the relative read counts of various time points. No upward trend in adapter efficiency was noted (Figure S1), indicating that a 2 minute incubation time is sufficient to create a library of sufficient quality to sequence and rapidly diagnose hotspot mutations.

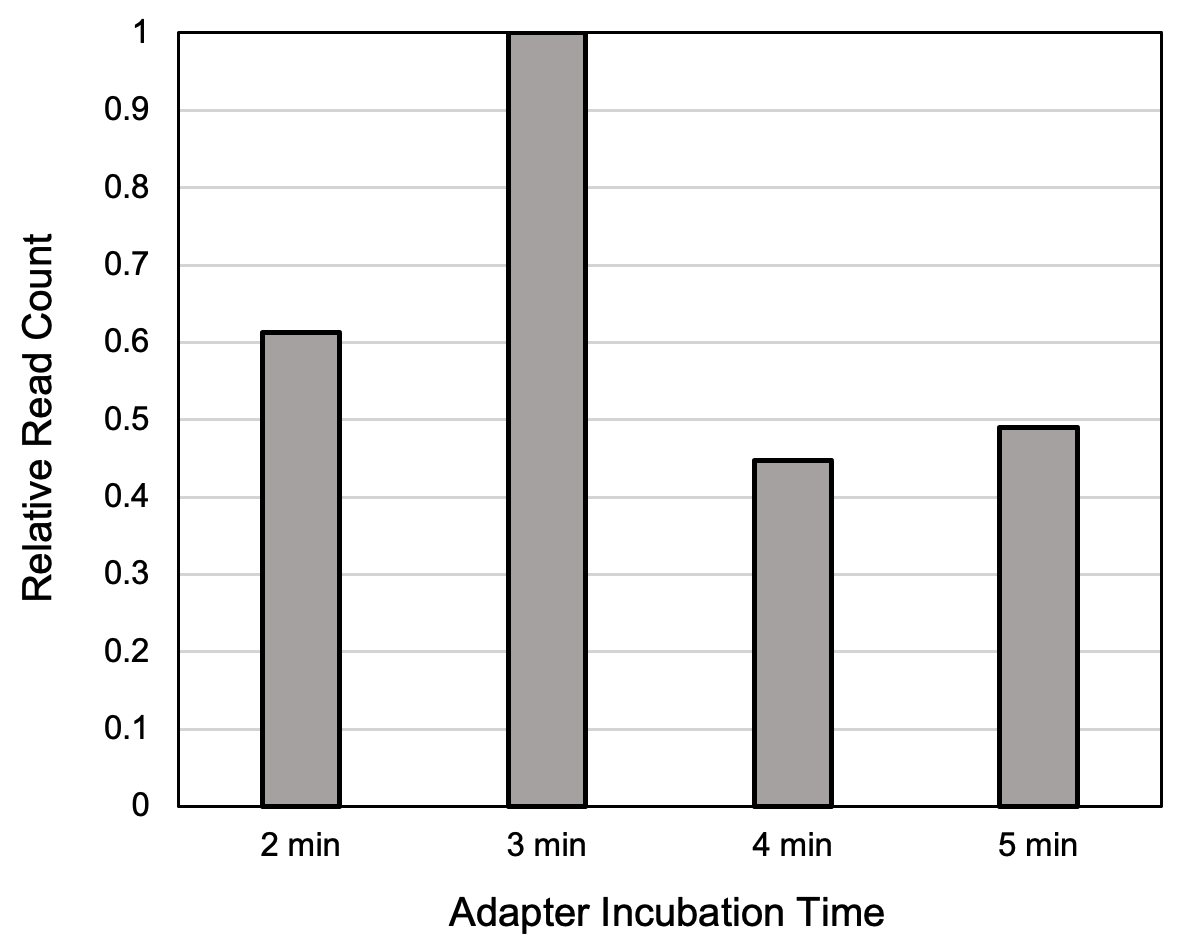

Figure S1. **Relative read counts from four different Rapid Adapter (RAP) incubation times.**

**LQE Dilution Efficiency and MinION Sequencing Feasibility**

To test the impacts on dilutions of the Lucigen QuickExtract (LQE) extracted DNA on PCR efficiency, two primer sets were designed to target exon 5 of the porcine (sus scrofa) TP53 gene. We used representative aliquots (~20mg) of pig brain tissue and followed the standard extraction protocol. Serial dilutions of extract were made and amplified using the following cycling parameters: 30s @ 98°C; then 35 cycles of 10s @ 98°C, 10s @ 62°C, 20s @ 72°C; then a final extension of 2 minutes @ 72°C. Amplification efficiency was measured via inspection of band brightness following gel electrophoresis (Figure S2). Dilutions of 8x-32x were substantially more efficient than 1x-4x with a 10x dilution (suggested by the manufacturer) performing the best across both primer sets.

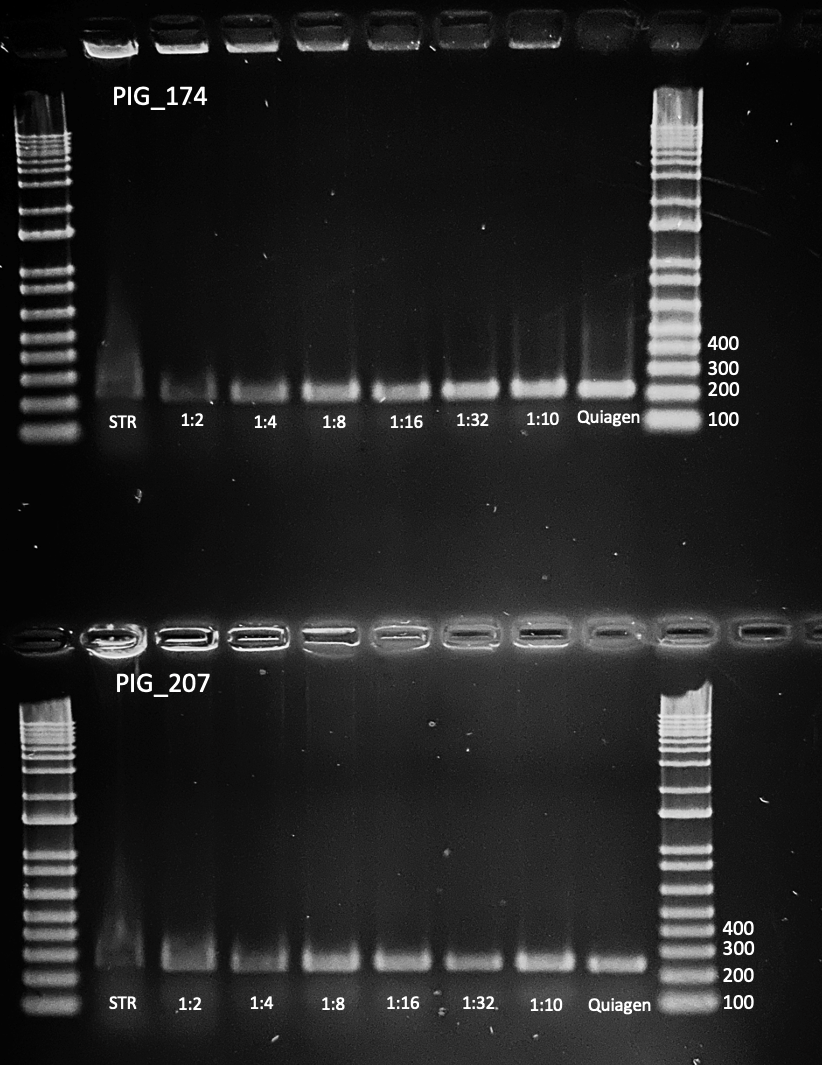

Figure S2. **Gel electrophoresis of PCR product from various dilutions of DNA extracted from Lucigen QuickExtract DNA extraction solution.** 10x (1:10) dilution performed best across both trials and was selected as a standard dilution for all further experiments.

We then performed an extraction on an aliquot of pediatric DIPG tumor and normal brain tissue following the standard LQE protocol and performed PCR with 10ul of 10x diluted DNA (1ul LQE product in 9ul nuclease free water). After confirming amplification, we sequenced product from 26 cycles of PCR to confirm that this product was sequenceable and produced valid reads and identified the known *HIST1H3B K27M* variant. We were able to successfully sequence and confirm the variant using our standard informatics pipelines, basecalling using ONTs Guppy basecaller and aligning reads to the human reference using minimap2 version 2.17. A read pileup visualized using integrated genome viewer is shown below.

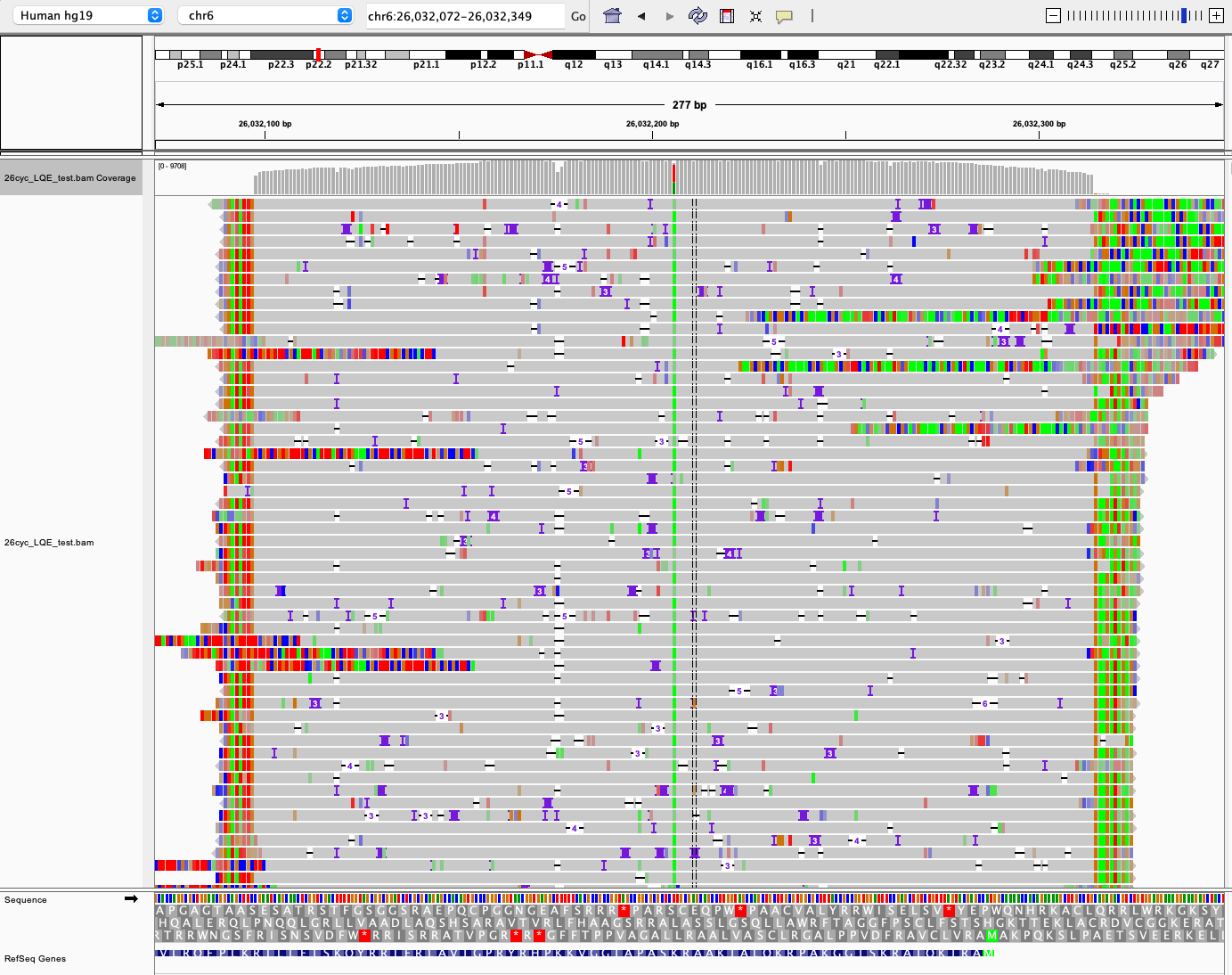

Figure S3. **Visualization of read pileup generated by Integrated Genome Viewer (IGV 2.8.0) showing fragmented amplicons aligned to *HIST1H3B.* The somatic *K27M* mutation is visible near the center of the alignment*.***

**LQE Incubation Time Efficiency**

DNA from prior standard protocol extractions was compared to DNA extracted from a separate aliquot of tumor tissue from the same patient. Extracted DNA was amplified using our primary *H3F3A K27M* LAMP assay in a Eppendorf Mastercycler, prepared using the suggested amount of dye. Amplification and fluorescence monitoring was performed at 65°C on an Eppendorf Mastercycler with 15s measurement intervals (cycles). Fluorescence curves are shown in Figure S4. The assay showed almost identical performance between the two extraction methodologies. Note that due to the inability to pre-heat the real-time fluorescence machine, and the difficulty of identifying the ratio of background DNA to the amplifying target, time-to-amplification and the sequencing threshold seems to be underestimated compared to our ultra-rapid protocol determined by time-sweep sequencing.

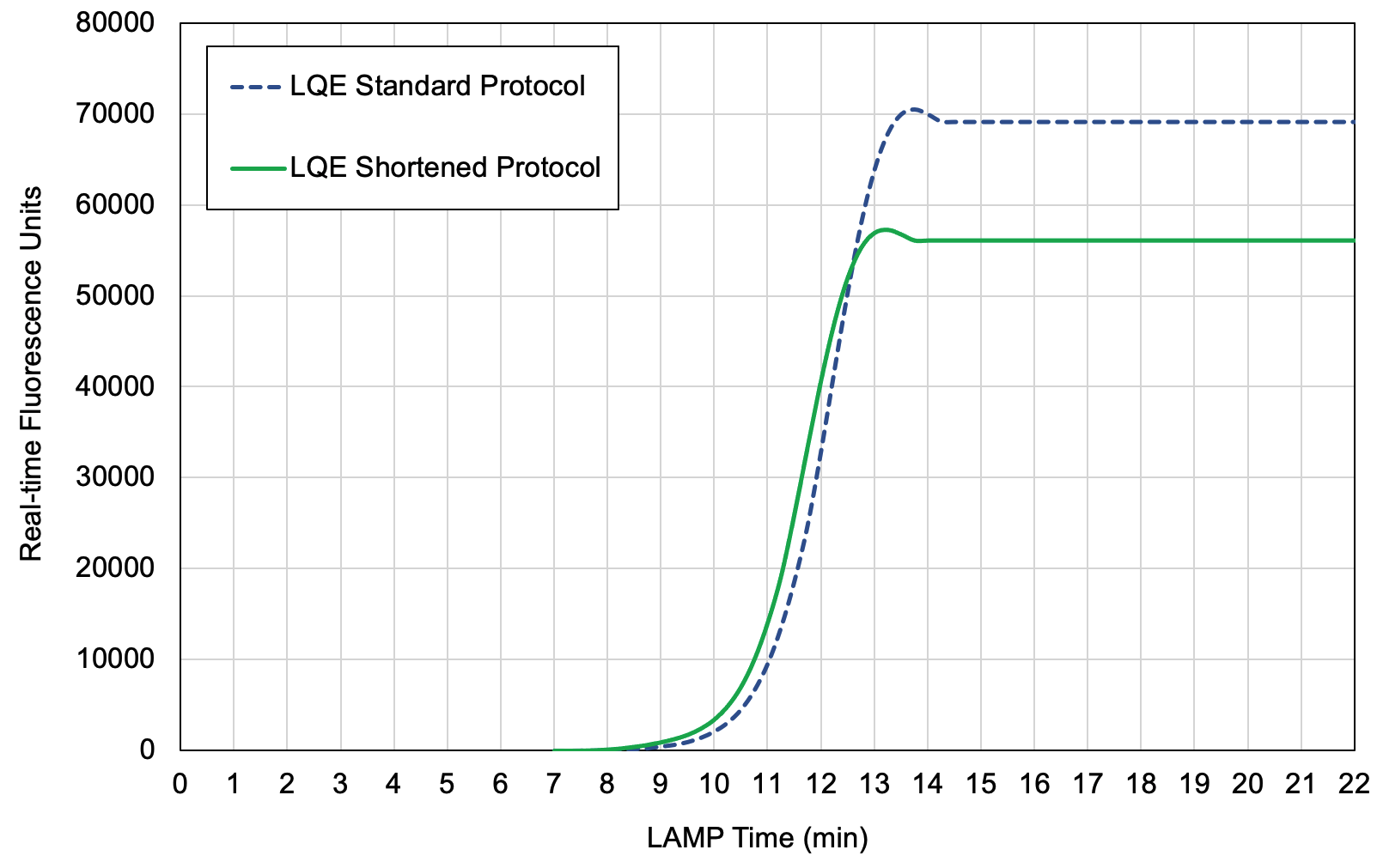

Figure S4. **Real-time amplification curves of DNA extracted using the standard LQE protocol and the shortened version.** The standard protocol leverages a 6 minute incubation time and the shortened protocol leverages a 1 minute incubation time. Amplification between the two methods is almost identical indicating that a reduced extraction time does not impact amplification rate, either due to reduced inhibitor concentration or unnecessarily long incubation time for this tissue type.

**Statistical Methods for Statistical Significance of Variant Call**

For our timed evaluations of the ultra-rapid sequencing protocols, we ended the protocol once read support over the target locus (mutant + wildtype calls) reached 250x. This is an arbitrary threshold and might not be enough coverage to call some very low variant allele fractions. In practice, *H3 K27M* fractions from tumor samples are high (20-60%). In this case, 250x is more than enough to call the variant with 99.99% confidence.

Statistical significance of a variant can be identified in real time, allowing Threshold Sequencing to be applied without an arbitrary coverage requirement. Our methodology involves computing the confidence interval for a proportion for each alternate variant call, or a targeted variant in the case of our *H3K27M* assay. The formula for computing the confidence interval is shown below:

$$\hat{p}\pm z\cdot\sqrt{\frac{\hat{p}(1-\hat{p})}{n}}$$

Where $\hat{p}$ is the observed variant proportion, $n$ is the target support (mutant and wild-type calls), and $z$ is the Z-value for the desired confidence level.

Given that ONT sequencers and corresponding basecallers have relatively high error rates, this confidence interval can be compared to the confidence interval for the expected error rate for that context. We call a variant statistically significant when the lower-bound confidence interval for the observed variant proportion exceeds the upper-bound confidence interval for the expected error rate proportion. We estimate the error rate for this locus to be ~1.5% based on prior characterization. Sequencing and basecaller errors are highly dependent on oligonucleotide sequence context and are much higher in low-complexity regions featuring homopolymers. This context-dependent error rate should be considered when calling variants in such regions. An example for a VAF proportion of 10% is shown in Figure S5. Alternatively, a healthy, wildtype sample can be pre-sequenced to characterize the error rate for a particular tissue, assay, target locus, and basecaller version. Future basecalling accuracy improvements will help reduce this error and lower sequencing depth requirements.

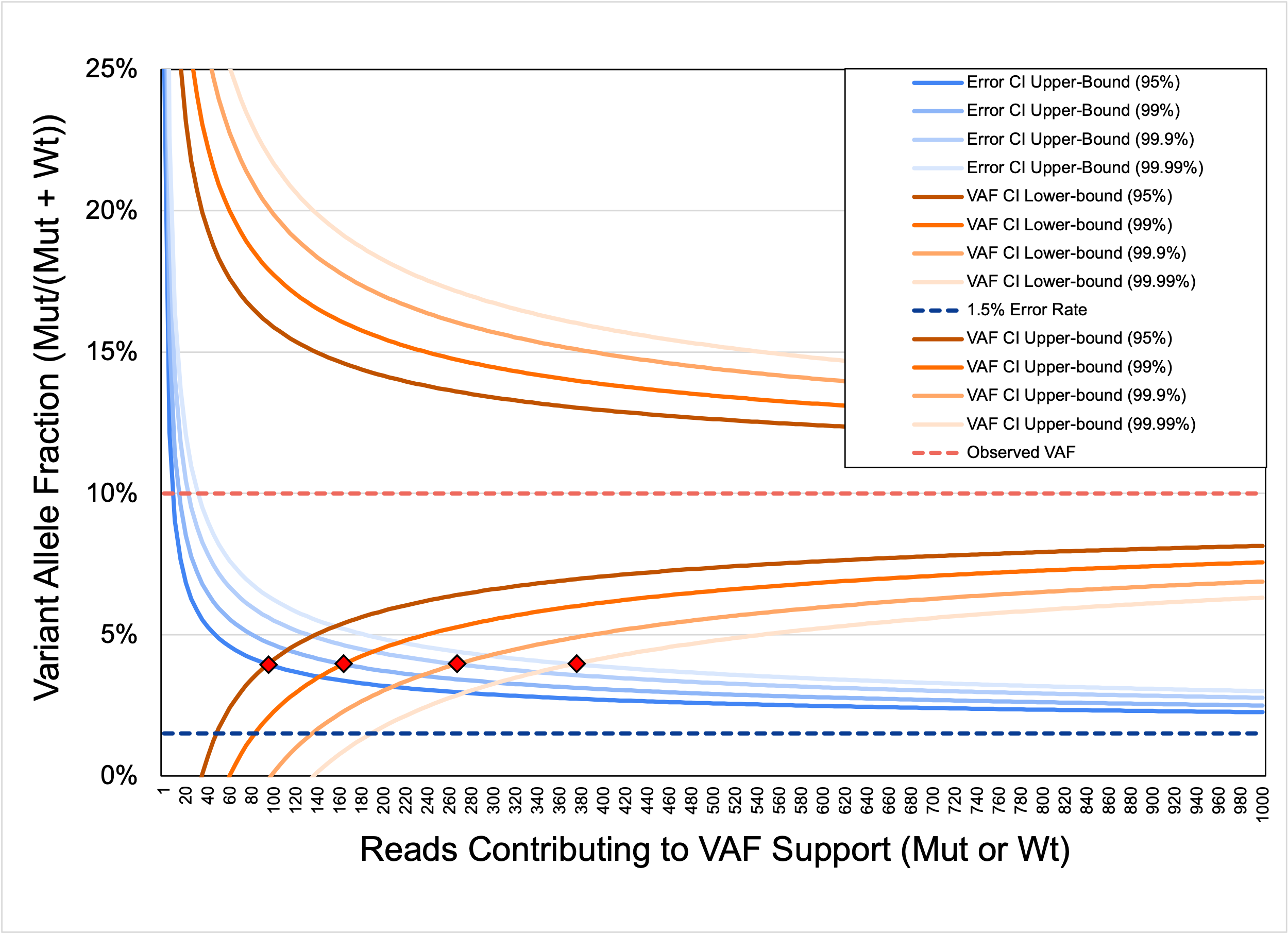

Figure S5. **Confidence intervals for simulated observed VAF of 10% (orange) relative to the confidence interval for estimated background 1.5% error rate (blue) plotted for various desired confidence levels (95%, 99%, 99.9%, 99.99%).** For each confidence level, we identify a variant call as statistically significant when the lower bound confidence interval of the observed variant fraction becomes larger than the upper bound confidence interval of the background error (red diamonds). Underlying VAF, error rate for that context, and desired confidence level greatly impact the locus support required to call a variant. In practice, it is best to apply these tests in real time. Even for a relatively low VAF of 10%, ~270 reads is expected to be sufficient to call the variant with 99.9% confidence.

**PCR Parameter Optimization and Efficiency Evaluation**

Faster PCR cycle parameters can lead to significant reductions in end-to-end amplification time. Using the best performing primer set evaluated (216bp) we performed cycle parameter optimization, reducing cycle times^12^ while balancing reaction efficiency. Due to the relatively small amplicon size allowed by the ONT rapid library preparation chemistry, PCR cycle times were able to be significantly reduced versus prior work^10^ but did end up relatively close to suggested lower-bound times (Figure S6a). Estimated end-to-end PCR times were modeled based on 3C/s ramp rate (Figure S6b). PCR paper time ignores ramp rate penalty and any handling penalties. Optimized PCR time is dominated by ramp rate of standard lab thermocyclers, and could be further reduced by using more rapid thermocyclers.

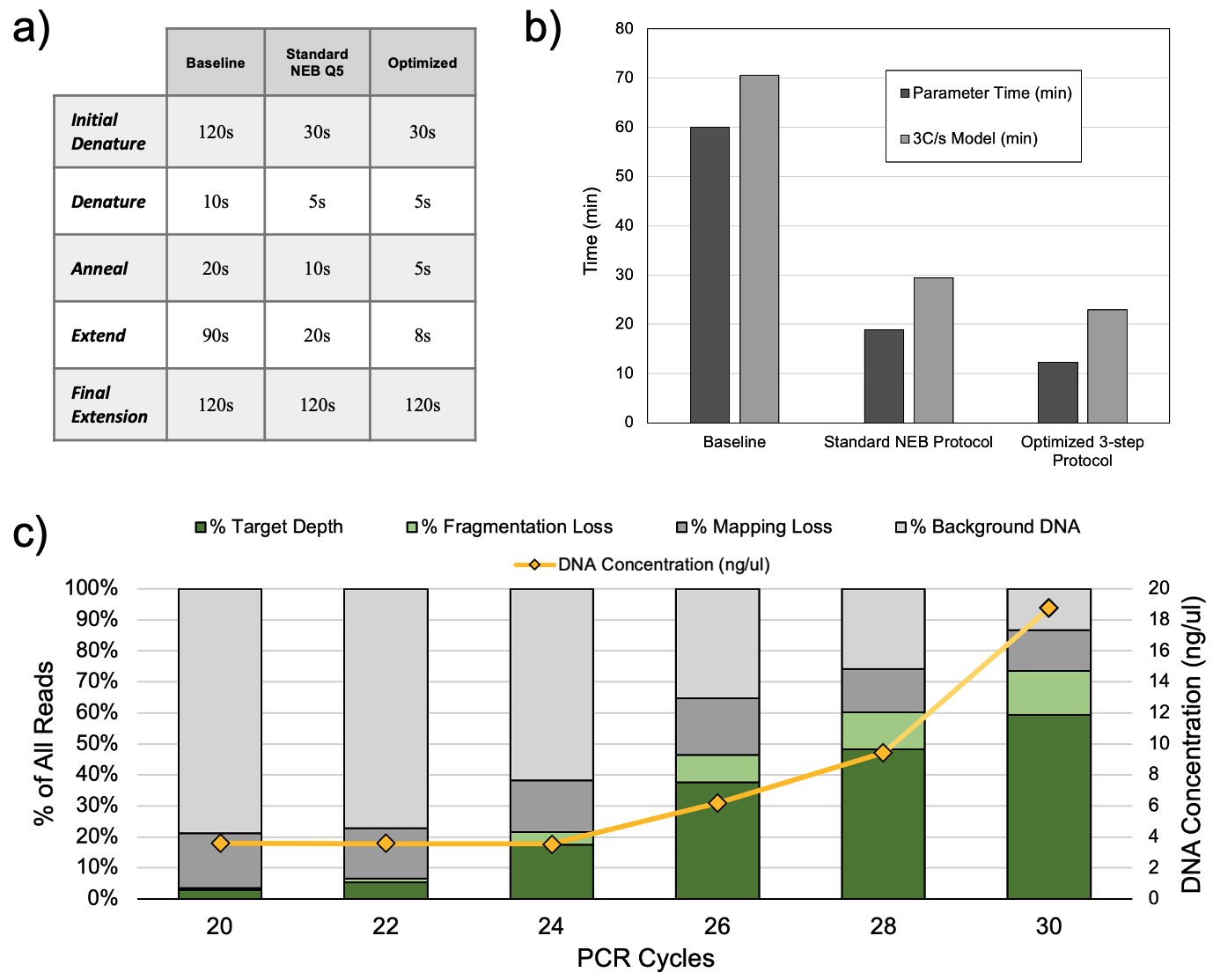

Figure S6. **3-step PCR assay optimized parameters, comparison to baseline protocols, and performance over various cycle counts. (a)** cycle times for a baseline amplification protocol^10^, the standard NEB protocol, and our our optimized parameters. **(b)** on-paper times differ significantly from modeled and measured times during experimentation. As cycle parameters are reduced, thermocycler ramp time begins to dominate total amplification time leading to a >50% efficiency loss (~10min penalty) and a reduction in the marginal benefit of cycle time reduction. **(c)** PCR performed well, producing a high-proportion of on-target amplicons after 26 cycles.

The proper cycle threshold for optimal end-to-end diagnostic performance can be estimated by sequencing product from PCR with varying numbers of cycles. We performed PCR using 1:10 diluted LQE extracted DNA (see Methods section). We then measured both the resulting PCR product concentration via Qubit fluorometer (Invitrogen; HS dsDNA Assay; #Q33230) and sequenced each resulting product using a barcoded, multiplexed rapid library preparation methodology (ONT; SQK-RBK004). Resulting reads were aligned to the human reference and classified according to alignment (see Methods section). Over time, we see both the total mass, as well as the proportion of PCR product relative to background genomic reads grow (Figure S6c).

**End-to-End PCR-based Threshold Sequencing Protocol**

The final end-to-end protocol is available in updateable electronic format at protocols.io (URL). A visual representation approximating the time-frame of the protocol as performed in this work is shown in Figure S7 along with a step-by-step protocol.

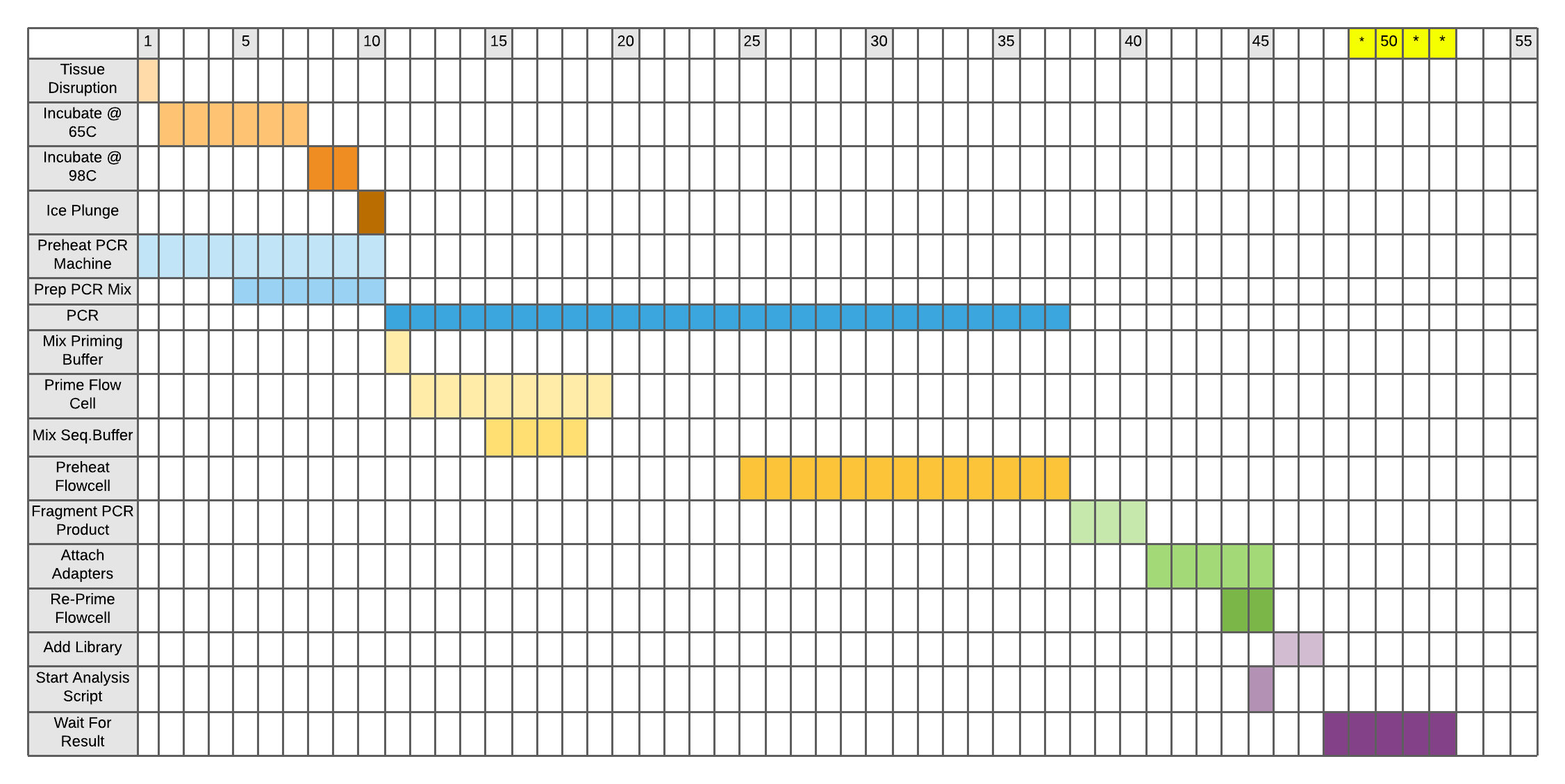

Figure S7. **Approximate protocol timeline showing time-frames and overlap of certain steps.**

**Ultra-rapid *HIST1H3B* PCR-based Assay Protocol**

1. **Prep**
   a. Pre-mix PCR mix according to 25ul Q5 2x Master Mix protocol adding 9ul of nuclease free water
   b. Heat two heat blocks to 65°C and 98°C respectively
   c. Pre-heat thermocycler
2. **DNA Extraction**
   a. Place ~20mg tissue in 500ul of Lucigen QuickExtract in a 2ml Eppendorf tube.
   b. Vortex for 10s
   c. Place tube in heat block at 65°C for 6 minutes
      i. Remove and briefly vortex on high after 3 minutes
   d. Place tube in heat block at 98°C for 2 minutes
   e. Plunge in ice for 30s
   f. Briefly vortex and spin down
3. **PCR**
   a. Extract 1ul of extracted DNA being careful not to clog the tip with any remaining viscous material or tissue
   b. Add 1ul of extracted DNA to PCR mix
   c. Flick to mix
   d. Spin down
   e. Insert into pre-heated thermocycler and initiate PCR program
   f. **Prep:**
      i. While PCR is running, mix the ONT priming mix and load into the flow cell
      ii. Initiate a sequencing run in the MinKNOW software (flow cell pre-heat and QC)
      iii. Pause the run once sequencing begins
4. **Library Preparation**
   a. Spin down PCR product
   b. Take 7.5ul of PCR product and add to 2.5ul ONT fragmentation buffer (FRA)
   c. Flick to mix
   d. Spin down
   e. Put back in thermocycler at 30°C for ONT fragmentation protocol
   f. Let thermocycler bring product down to room temp
   g. Add 1ul of ONT rapid adapter (RAP) to tube
   h. Flick to mix
   i. Incubate at room temperature for 5 minutes (flicking and spinning down if desired)
5. **Loading and Sequencing**
   a. With 1 minute remaining in RAP incubation, re-prime MinION flow cell
   b. Mix entire library into sequencing mix, making sure to mix by pipetting to ensure library contact with loading beads
   c. Drop all 75ul of sequencing library into the MinION SpotOn port and unpause sequencing run

**2-step PCR and Four-primer Protocol Evaluation**

Two-step PCR combines the annealing and extension steps, reducing both ramp penalty and generally overall cycle time. We explored design and evaluation of a 2-step protocol to further reduce the time cost of PCR-based target amplification. The 2-step evaluation was successful, but not evaluated in and end-to-end setting do to resource constraints and the rapid amplification rate of LAMP.

Oxford Nanopore’s Four Primer Protocol (ONT; SQK-PSK004) involves a two stage amplification. The first stage uses target specific primers tailed with ONT-specific handshake sequences to amplify and tail the target of interest. The second stage uses ONT provided primers that contain just the handshake sequences, tailed with click chemistry. This allows for fragmentation free library preparation, obviating both the time penalty of the fragmentation step and the fragmentation penalty. In practice, we found this protocol to be difficult to optimize and perform due to its two-separate amplification stages and not worth the optimization time for the end-to-end time benefit especially given the rapid amplification rate of LAMP. However, we did develop a single stage protocol that successfully amplified and tailed primer sequences. We found that aggressively reducing cycle times negatively impacted efficiency and negated the time-improvement of the protocol. However, we report our progress here for posterity and future optimization efforts. All steps should be followed according to ONT’s SQK-PSK004 or SQK-PBK004 protocols except for the amplification step which should be replaced by the following

1. Mix PCR reaction as follows
   a. 12.5ul NEB Q5 2x Hot Start Master Mix (NEB #M0494)
   b. 1.25ul 1uM 216 Fwd Primer (50nM final concentration)
   c. 1.25ul 1uM 216 Reverse Primer (50nM final concentration)
   d. 1.5ul ONT WGP (SQK-PSK004) diluted 1:2 in nuclease free water
   e. 8.5ul Qiagen DNAEasy (Qiagen #69504) extracted DNA (27.16ng/ul in our experiment)
2. Perform PCR according to cycling parameters in Table S2
   a. 30s @ 98°C
   b. 35 cycles of
      i. 5s @ 98°C
      ii. 10s @ 64°C
      iii. 13s @ 72°C
   c. 30s @ 72°C
3. Sequence according to standard ONT SQK-PSK004 or SQK-PBK004 protocols

**LAMPrey Algorithm Description**

LAMPrey is an algorithm for processing a set of FASTQ reads, attempting to identify properly formed LAMP concatemers (lamplicons) and call variants based off of a pileup of lamplicon sub-reads. As input, the user supplies a FASTQ file of sequenced LAMP product and a corresponding primer sequence file that details the 6-8 design sequences identified in LAMP primer design (B3, B2, B1, F1, F2, F3, BLP, FLP), as well as other sequences of interest such as ONT barcodes or adapter sequences. The user also describes a “target” sequence that spans the location where the mutation of interest lies. The user also specifies where this target sequence lies in the LAMP design sequence schema. A target sequence can exist anywhere between the F2 and B2 primers as long as it is not covered by another primer.

**Sequence Alignment:** LAMPrey first parses the FASTQ read file and optionally extracts the reads according to the ONT Guppy basecaller emitted *start_time* timestamp. This is assumed to be the approximate order and time that the reads are generated. LAMPrey then considers each read and aligns each sequence in the primer file to the read using a standard Smith Waterman alignment. Alignments are recorded if their identity is above a certain threshold (default of 75%). Once all possible primers are marked, overlapping aligned sequences are removed, prioritizing alignments with higher identity. After sequence alignment, reads can be immediately classified into one of five different categories.

**Classification:** If the read does not contain any aligned target sequences, it is diagnosed as either an amplicon fragment, background genomic DNA, spurious LAMP amplification, an ONT fragment leftover from library preparation, or Unknown (further broken down into “short” unknown sequences (<60bp) and other Unknown). Amplicon fragments are those that align to the gene or target region of interest within some expected distance (e.g. 1000bp) but do not contain the target sequence. These reads are expected given the fragmentation step included in library preparation. If the read aligns somewhere else in the human genome, it is marked as a background genomic read. These are also expected given that Threshold Sequencing skips target enrichment to save diagnostic time. If a read has a large proportion of primer sequences covering the read, but does not contain a target sequence, it is marked as suspected spurious amplification. These are assumed to be hybridized primers or other undesired amplification that does not capture the diagnostically relevant information of the patient, but still involves the basic LAMP machinery and at least some primer sequences. These reads are not expected but are difficult to fully avoid without extensive optimization of the LAMP assay. If a read has a high proportion of ONT related sequences covering a large proportion of the read, it is marked as an ONT fragment. ONT fragments could be unligated adapter sequences, or read fragments that contain too little target or background DNA to successfully align to the reference. If a read cannot be classified as any of the other categories, it is marked as unknown, with reads shorter than 60bp marked as “short”.

**Chaining:** Once a lamplicon with at least one target sequence is identified, it is further processed to divide up the possible concatemer into its sub-reads. This is accomplished by considering each target sequence and looking for expected primer sequences to the 5’ and 3’ ends based on the LAMP assay. This is analogous to the chaining of read hits in many common read mapping algorithms. For example, if a forward target sequence exists between the F1 and B1 regions, we would look to its 5’ end to find an aligned F1 sequence, and to its 3’ end to find the complement of the B1 sequence. In this way, target sequences are “extended” in the 5’ and 3’ direction until an aligned sequence that doesn’t match the expected ordering is found (e.g. an ONT barcode, adapter sequence, or FIP/BIP loop transition), or the end of the read is found. This extension defines a sub-read. Each sub read is extracted and considered separately.

**Variant Calling:** Once each sub-read is identified, they are separately aligned to the target gene of interest (NOTE: not the full human reference). All sequences that successfully align are converted to a pileup, and the base pairs over the target hotspot locus are considered. Because LAMP concatemers contain N target copies, and ONT sequencers and basecalling algorithms may introduce errors, the N target copies may not agree. For variant calling, we only consider lamplicons that have majority or plurality support for one call from all alignable sub-reads. For example, a lamplicon concatemer with three sub-reads aligned over the target locus with calls CCT would be considered an C call (majority and plurality). A concatemer with two target regions with calls CT would be ignored (majority) or a random basecall would be selected from the plural basecalls (plurality). Currently, variant allele fraction is computed in real time as reads are processed based off of a user supplied VCF file describing a single variant of interest, however, LAMPrey could be easily configured to identify arbitrary mutations at multiple loci.

**End-to-End LAMP-based Threshold Sequencing Protocol**

The final end-to-end protocol is available in updateable electronic format at protocols.io (URL). A visual representation approximating the time-frame of the protocol as performed in this work is shown in Figure S8 along with a step-by-step protocol. This final protocol combines DNA extraction, amplification, and library preparation into one thermocycler program to save time and also simplify equipment usage. The program is shown below with pause points.

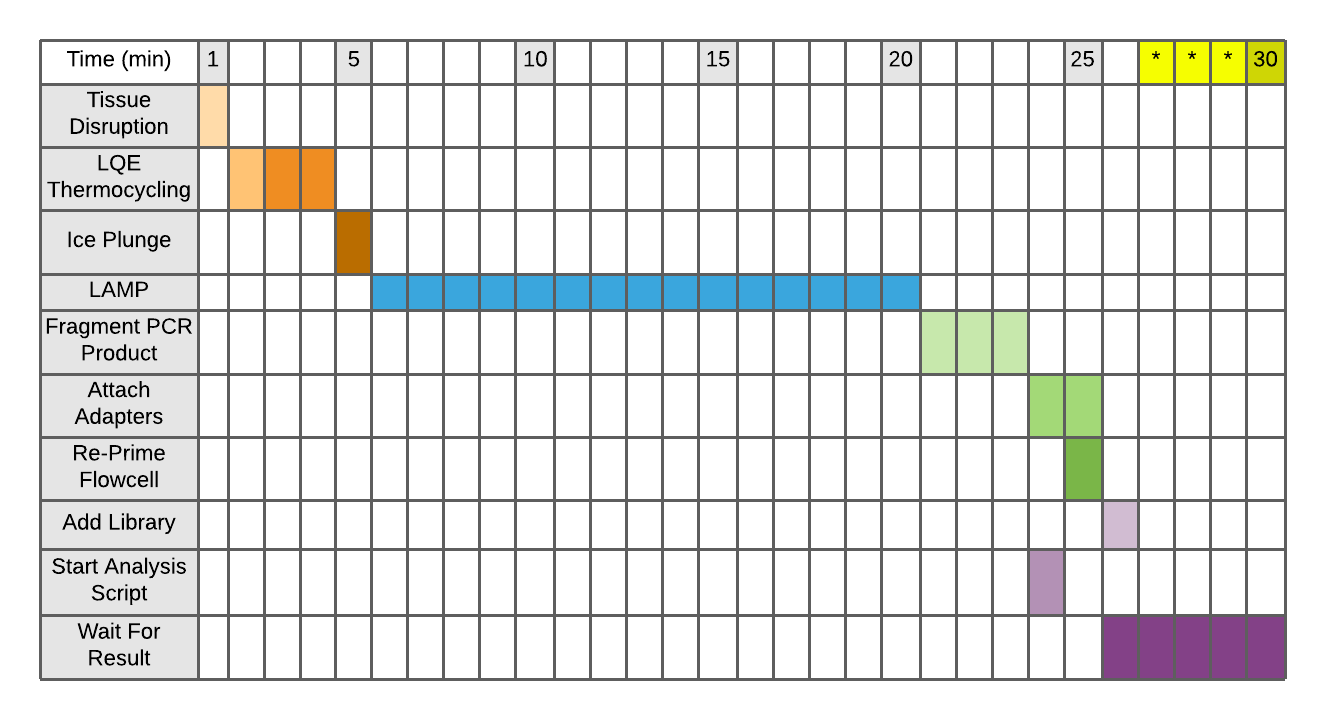

Figure S8. **Steps and time-frame of performed end-to-end LAMP-based diagnostic.** Yellow highlighted region delineates the time-frame of an expected result. We achieved a variant call at ~29.5 minutes. Flow cell priming, LAMP mix preparation, flow cell QC, sequencing buffer mixing, and fragmentation buffer preparation was all performed ahead of time.

**Ultra-rapid H3F3A LAMP-based Assay Protocol**

1. **Prep**
   a. Prime flow cell according to ONT SQK-RAD004 protocol and wait five minutes
   b. Initiate a sequencing run in the MinKNOW software (flow cell pre-heat and QC)
   c. Pause the sequencing run once sequencing begins
   d. Pre-mix LAMP reaction mix according to 25ul NEB WarmStart LAMP 2x Master Mix protocol in a 0.2ml PCR tube
      i. 12.5ul LAMP 2x WarmStart Master Mix
      ii. 2.5ul 1x LAMP primer mix
      iii. 9ul Nuclease free water
   e. Pre-mix fragmentation mix in a 0.2ml PCR tube
      i. 2.5ul ONT FRA
      ii. 5.6ul nuclease free water (to account for a 4x dilution of LAMP product)
   f. Pre-mix sequencing mix in a 0.2ml PCR tube according to ONT SQK-RAD004 protocol and place on ice
   g. Pre-heat thermocycler
2. **DNA Extraction**
   a. Place ~20mg tissue in 100ul of Lucigen QuickExtract in a 0.2ml PCR tube
   b. Vortex for 30s and spin down
   c. Place tube in thermocycler and initiate program
   d. Pause program after LQE deactivation portion has finished
   e. Plunge in ice for 10s to bring to room temp
   f. Vortex on high briefly and spin down
3. **LAMP**
   a. Extract 1ul of extracted DNA being careful not to clog the tip with tissue
   b. Add 1ul of extracted DNA to LAMP mix
   c. Flick to mix and spin down
   d. Insert into thermocycler and unpause program
   e. Pause thermocycler program after LAMP incubation portion has finished
4. **Library Preparation**
   a. Spin down LAMP product
   b. Add 1.9ul to fragmentation mix
   c. Flick to mix
   d. Spin down
   e. Insert into thermocycler and unpause program
   f. Plunge in ice for 10s
   g. Add 1ul of ONT rapid adapter (RAP) to tube
   h. Flick to mix and spin down
   i. Incubate at room temperature for 2 minutes
5. **Loading and Sequencing**
   a. With 1 minute remaining in RAP incubation, re-prime MinION flow cell with SpotOn port open
   b. Mix entire 11ul library into sequencing mix, making sure to mix by pipetting to ensure library contact with loading beads
   c. Drop all 75ul of sequencing library into the MinION SpotOn port and unpause sequencing run

**References**

1.	Terata, K. *et al.* Novel rapid-immunohistochemistry using an alternating current electric field for intraoperative diagnosis of sentinel lymph nodes in breast cancer. *Sci. Rep.* **7**, 2810 (2017).

2.	Lovchik, R. D., Taylor, D. & Kaigala, G. Rapid micro-immunohistochemistry. *Microsyst. Nanoeng.* **6**, 1–10 (2020).

3.	Duncan, D. J., Vandenberghe, M. E., Scott, M. L. J. & Barker, C. Fast fluorescence in situ hybridisation for the enhanced detection of MET in non-small cell lung cancer. *PLOS ONE* **14**, e0223926 (2019).

4.	Livermore, L. J. *et al.* Rapid intraoperative molecular genetic classification of gliomas using Raman spectroscopy. *Neuro-Oncol. Adv.* **1**, vdz008 (2019).

5.	Al-Ramadhani, S. *et al.* Metasin—An Intra-Operative RT-qPCR Assay to Detect Metastatic Breast Cancer in Sentinel Lymph Nodes. *Int. J. Mol. Sci.* **14**, 12931–12952 (2013).

6.	Smith, G. J., Hodges, E., Markham, H., Zhang, S. & Cutress, R. I. Evaluation of the Metasin assay for intraoperative assessment of sentinel lymph node metastases in breast cancer. *J. Clin. Pathol.* **70**, 134–139 (2017).

7.	Kanamori, M. *et al.* Rapid and sensitive intraoperative detection of mutations in the isocitrate dehydrogenase 1 and 2 genes during surgery for glioma. *J. Neurosurg.* **120**, 1288–1297 (2014).

8.	Gm, S. *et al.* Rapid Intraoperative Molecular Characterization of Glioma. *JAMA Oncol.* **1**, 662–667 (2015).

9.	Bamborschke, D. *et al.* Ultra-rapid emergency genomic diagnosis of Donahue syndrome in a preterm infant within 17 hours. *Am. J. Med. Genet. A.* **n/a**,.

10.	Euskirchen, P. *et al.* Same-day genomic and epigenomic diagnosis of brain tumors using real-time nanopore sequencing. *Acta Neuropathol. (Berl.)* **134**, 691–703 (2017).

11.	Minimap2: pairwise alignment for nucleotide sequences | Bioinformatics | Oxford Academic. https://academic.oup.com/bioinformatics/article/34/18/3094/4994778.

12.	Protocol for Q5® Hot Start High-Fidelity 2X Master Mix | NEB. https://www.neb.com/protocols/2012/08/30/protocol-for-q5-hot-start-high-fidelity-2x-master-mix-m0494.
